# The molecular architecture of the desmosomal outer dense plaque by integrative structural modeling

**DOI:** 10.1101/2023.06.13.544884

**Authors:** Satwik Pasani, Kavya S Menon, Shruthi Viswanath

**Affiliations:** National Center for Biological Sciences, Tata Institute of Fundamental Research, Bengaluru 560065, India

**Keywords:** integrative structural modeling, macromolecular assemblies, cell-cell junctions, desmosome

## Abstract

Desmosomes mediate cell-cell adhesion and are prevalent in tissues under mechanical stress. However, their detailed structural characterization is not available. Here, we characterized the molecular architecture of the desmosomal outer dense plaque (ODP) using Bayesian integrative structural modeling *via* the Integrative Modeling Platform. Starting principally from the structural interpretation of an electron cryo-tomogram, we integrated information from X-ray crystallography, an immuno-electron microscopy study, biochemical assays, *in-silico* predictions of transmembrane and disordered regions, homology modeling, and stereochemistry information. The integrative structure was validated by information from imaging, tomography, and biochemical studies that were not used in modeling. The ODP resembles a densely packed cylinder with a PKP layer and a PG layer; the desmosomal cadherins and PKP span these two layers. Our integrative approach allowed us to localize disordered regions, such as N-PKP and PG-C. We refined previous protein-protein interactions between desmosomal proteins and provided possible structural hypotheses for defective cell-cell adhesion in several diseases by mapping disease-related mutations on the structure. Finally, we point to features of the structure that could confer resilience to mechanical stress. Our model provides a basis for generating experimentally verifiable hypotheses on the structure and function of desmosomal proteins in normal and disease states.

**Significance statement:** Desmosomes are cell-cell junctions that possess a hyper-adhesive property and are prevalent in tissues under mechanical stress. However, their detailed structural characterization has eluded experimental structural biologists so far. Here, we use an integrative approach that allows us to rigorously combine biochemical, biophysical, and cell biological data at multiple scales in order to determine the molecular architecture of the outer dense plaque region of desmosomes. We validate the structural model by several pieces of information not used to compute it. The model allows us to generate hypotheses on the desmosomal proteins in normal and disease states.

## Introduction

Desmosomes are large, 300nm-long protein assemblies that connect the keratin intermediate filaments of adjacent cells. They mediate cell-cell adhesion and play a crucial role in maintaining tissue integrity for tissues under mechanical stress, such as heart and epithelial tissues. They also play critical roles in cell signalling and tissue differentiation. Dysfunction of desmosomes has been implicated in skin and heart diseases, auto-immune diseases, and cancers (Garrod & Chidgey, 2008; Green & Simpson, 2007; Kowalczyk & Green, 2013).

The ultra-structure of desmosomes shows its organization in three areas: the extracellular core region (EC), the outer dense plaque (ODP), and the inner dense plaque (IDP)(Delva et al., 2009). The EC is made up of the desmosomal cadherins (DCs), desmoglein (DSG) and desmocollin (DSC), which interact with similar molecules in adjacent cells to achieve cell-cell adhesion. The ODP, which spans 15-20nm, is a protein dense region between the EC and IDP. Here, members of the armadillo family - plakoglobin (PG) and plakophilin (PKP), members of the plakin family-desmoplakin (DP), and the cytoplasmic tails of the desmosomal cadherins interact. The ODP functions to regulate cadherins, since it contains several phosphorylation sites and binding sites for regulatory proteins (Badu-Nkansah & Lechler, 2020; Garrod & Chidgey, 2008). Recent proteomics studies have identified several regulatory proteins that are seen in the ODP (Badu-Nkansah & Lechler, 2020). Desmoplakin links to the keratin intermediate filaments in the IDP at the cytoplasmic end of the desmosome (Garrod & Chidgey, 2008; Kowalczyk & Green, 2013).

A detailed structural characterization of the ODP is not yet available. A molecular map based on immuno-electron microscopy is known (North et al., 1999). However, this map provides the distances of plaque protein termini from the plasma membrane; it does not provide information on the three-dimensional arrangement of the proteins. A 32Å cryo-electron tomogram of the ODP, which shows its organization in two layers, has been determined by (Al-Amoudi et al., 2011). This is also the most comprehensive structural study on the ODP so far. However, the resolution of the tomogram is too low to unambiguously fit the known structures of plaque proteins and protein complexes. Moreover, domains of unknown structure, comprising a significant portion of the ODP, were not modeled. These domains make up about 40% of the protein sequences of the stratified epithelial desmosomal ODP (Fig. 1A, Table S1). In this study, we built a more complete model of the ODP, including domains of unknown structure, by combining the data from electron cryo-tomography and immuno-EM experiments with an array of known biophysical, biochemical, and cell biological experimental data, bioinformatics predictions, and physical principles (Fig. 1, Table S2-S3) (Bonné et al., 2003; Bornslaeger et al., 2001; Hatzfeld et al., 2000; Kowalczyk et al., 1999; Smith & Fuchs, 1998).

**Figure 1.**
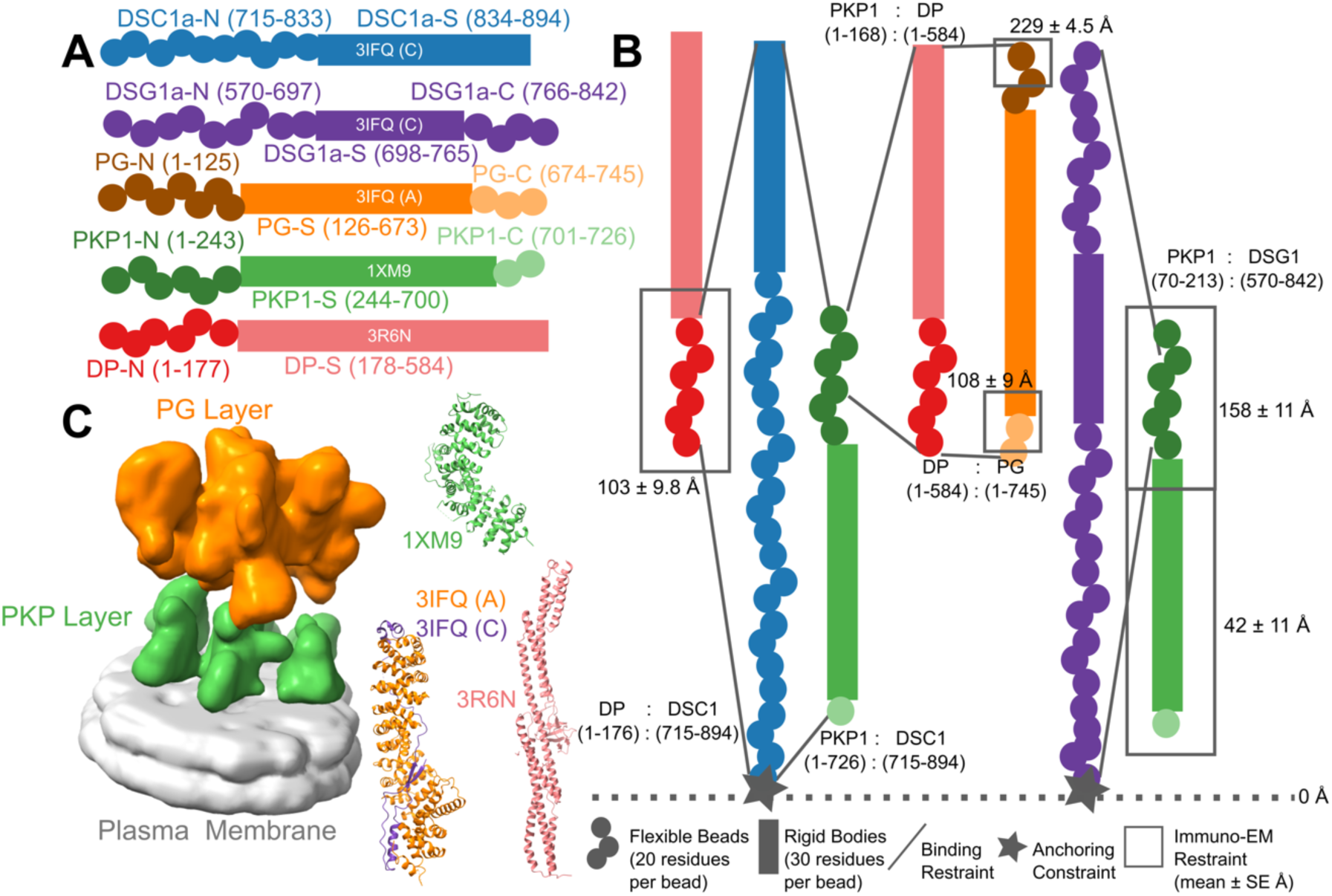
Representation and restraints used for integrative modeling of the desmosomal ODP. **A)** The isoforms used in modeling the desmosomal ODP of stratified epithelia and the representation of the different protein domains as rigid bodies with known structure (rectangles with PDB ID and chain name) or flexible beads (circles). The domains with known structure are usually denoted by a suffix -S after the protein (*e.g.,* DP-S), while the termini are denoted by -N or -C suffixes after the protein (*e.g.,* DP-N). **B)** Three types of restraints are shown. **1.** Binding restraints between interacting protein domains depicted by a pair of lines connecting the boundaries of each interacting domain pair. **2.** Immuno-EM restraint for localizing protein termini depicted by rectangles around the restrained protein terminus, and **3.** Anchoring constraint for localizing the transmembrane region of the cadherins depicted by star. The color scheme follows that in Panel A. **C) (Left)** The cryo-electron tomogram (EMD-1703) used for modeling with the PKP and the PG layers segmented. The density corresponding to the plasma membrane was not used for modeling. **(Right)** The PDB structures used, colored according to panel A. See also Methods, Fig. S1, Tables S1-S2.

Structures of large protein assemblies such as desmosomes are challenging to characterize using a single experimental method such as X-ray crystallography or cryo-electron microscopy. Purifying the component proteins is difficult since several of these are membrane proteins. Here we applied integrative structural modeling *via* IMP (Integrative Modeling Platform; https://integrativemodeling.org) to characterize the molecular architecture of the ODP (Alber et al., 2007; Rout & Sali, 2019; Russel et al., 2012). In this approach, we combined information from experiments along with physical principles, statistical inference, and prior models for structure determination. Several assemblies have been determined using this approach, including the yeast nuclear pore complex (Alber et al., 2007; Kim et al., 2018), 26S proteasome (Lasker et al., 2012), yeast centrosome (Viswanath, Bonomi, et al., 2017), and chromatin-modifying assemblies (Arvindekar et al., 2022; Robinson et al., 2015). Importantly, the Bayesian inference framework allowed us to rigorously and objectively combine multiple sources of experimental data at different spatial resolutions by accounting for the data uncertainty. It also facilitated the modeling of full-length proteins, including regions of unknown structure and/or disorder along with regions of known and/or readily modeled atomic structure.

Based primarily on the structural interpretation of the electron cryo-tomogram of the ODP (Al-Amoudi et al., 2011), we integrated information from an immuno-electron microscopy study, several X-ray crystallography studies, biochemical studies based on yeast two-hybrid, co-immunoprecipitation, *in vitro* overlay, and *in vivo* co-localization assays, *in-silico* sequence-based predictions of transmembrane and disordered regions, homology modeling, and stereochemistry information to obtain the integrative structure of the desmosomal ODP (Fig. 1-2, Table S1-S3). Our structure was further validated by additional information super-resolution imaging, newer tomograms, and biochemical studies not used in modeling (Choi et al., 2009; Kowalczyk et al., 1997; Smith & Fuchs, 1998; Troyanovsky, Troyanovsky, Eshkind, Krutovskikh, et al., 1994; Troyanovsky, Troyanovsky, Eshkind, Leube, et al., 1994; Wahl et al., 1996) (Table S3). Our approach allowed us to localize disordered regions such as the N-terminus of plakophilin and the C-terminus of plakoglobin in the context of regions of known structure.

**Figure 2.**
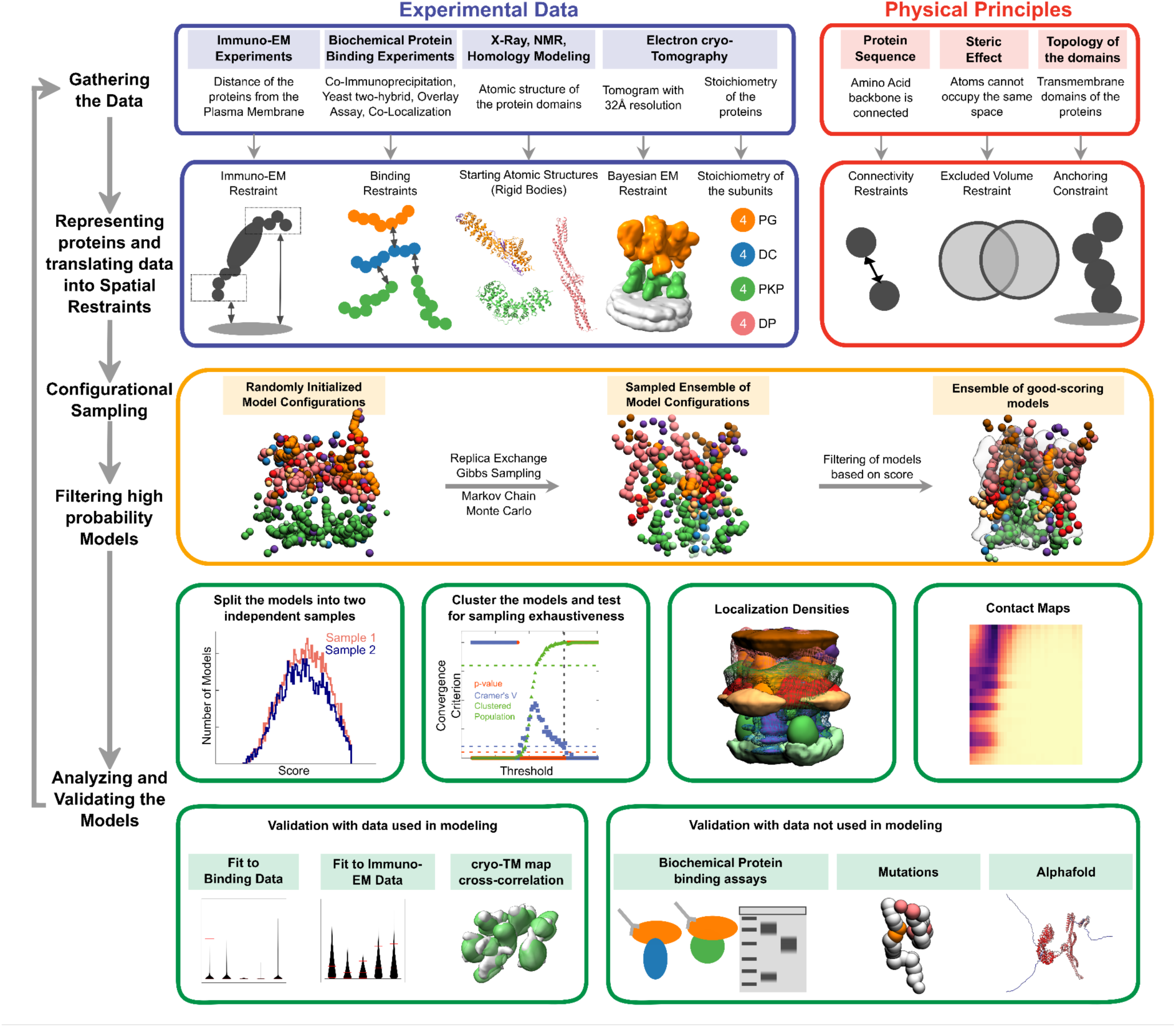
Integrative modeling of the desmosomal ODP. From top to bottom, the rows describe the input information (first), how the input information is encoded into spatial restraints (second), the sampling procedure (third), analysis (fourth), and validation of the results (fifth).

We refine known protein-protein interactions in the ODP, provide structure-based hypotheses for defective cell-cell adhesion associated with pathogenic mutations seen in skin diseases and cancers, and identify aspects of the desmosome structure that could possibly confer robustness to mechanical stress. Further, our integrative structure is more complete in terms of the sequence coverage of the ODP proteins compared to other structures, for *e.g.*, based on cryo-electron tomography, which lack the disordered domains of ODP proteins (Al-Amoudi et al., 2011) (*e.g.,* N-Pkp). The current work forms a basis for generating experimentally verifiable hypotheses on the structure and function of desmosomal proteins.

## Results and Discussion

### Summary of the integrative modeling workflow

The expression of isoforms of the ODP subunits is tissue-dependent (Delva et al., 2009; Green & Simpson, 2007). Below, we detail the integrative structure of the desmosomal ODP corresponding to the upper epidermis, comprising of plakoglobin (PG), desmoplakin (DP), plakophilin (PKP1), desmocollin (DSC1a, henceforth DSC1), and desmoglein (DSG1a, henceforth DSG1) (Fig. 1A, Table S1). The upper epidermis was chosen since the isoforms predominantly expressed in stratified epithelial tissue were associated with the most data. In contrast, very little data that can be used for integrative structural modeling, *e.g.*, biochemical interaction data, was available on the cardiac isoforms was available on the cardiac isoforms. For example, a single study reports on PKP2 biochemical interactions with other ODP proteins (Chen et al., 2002), whereas PKP1 biochemical interactions are studied in at least four papers (Table S2). Little to no protein-protein interaction data was found for cardiac isoforms DSG2/DSC2/DP2. Finally, the tomography data and immuno-EM data also correspond to epithelial tissue. As a simplifying assumption, our model of the ODP contains a single isoform of PKP, DSC, and DSG, corresponding to the dominant isoform in the modeled stratified epithelial tissue.

The protein domains constituting the desmosomal ODP and the corresponding terminology used henceforth are shown (Fig. 1A, Table S1). The stoichiometry of these proteins was determined using a previously published cryo-electron tomogram (Methods) (Al-Amoudi et al., 2011). Integrative modeling proceeded in four stages (Fig. 2, Methods). Data from X-ray crystallography, electron cryo-tomography, immuno-electron microscopy, and biochemical assays was integrated with *in-silico* sequence-based predictions of transmembrane and disordered regions, homology modeling, and stereochemistry information (Fig. 1B-1C, Table S2, Table S3).

Each protein was represented by a series of spherical beads along the backbone, each bead denoting a fixed number of residues. Protein domains with X-ray structures or homology models (such as the PKP1 armadillo repeat domain) were represented at 30 residues per bead and modeled as rigid bodies, whereas domains without known atomic structure (such as the PKP1-N) were coarse-grained at 20 residues per bead and modeled as flexible strings of beads (Fig. 1A, 1C, Table S1, Methods). Data from immuno-EM was used to restrain the distance of protein termini from the plasma membrane, cryo-electron tomograms were used to restrain the localization of ODP proteins, and the data from biochemical assays restrained the distance between interacting protein domains (Fig 1B, Methods). Starting with random initial configurations for the rigid bodies and flexible beads, 180 million models were sampled using Replica Exchange Gibbs Sampling MCMC, from a total of 50 independent runs. At each step, models were scored based on agreement with the immuno-EM, tomography, and biochemical data, together with additional stereochemical restraints such as cylinder restraints, connectivity, and excluded volume (see Methods).

About 24866 good-scoring models were selected for further analysis (see Methods, Stage 4 for details). These models were clustered based on structural similarity and the precision of the clusters was estimated (Arvindekar et al., 2022; Saltzberg et al., 2021; Viswanath, Chemmama, et al., 2017) (Fig. S2). The quality of the models was assessed by the fit to input data, as well as to data not used in modeling (Fig. S3-S4, Table S2-S3, Methods). Further analysis included identification of protein-protein interfaces *via* contact maps and rationalizing skin and cancer-related diseases involving ODP proteins *via* mapping of known missense, pathogenic mutations on the integrative structure (Fig. 4-5, Fig. S5, Table S4-S5, Methods).

### Integrative structure of the desmosomal ODP in the upper epidermis

Integrative modeling of the desmosomal ODP in the upper epidermis resulted in a single cluster of 24016 models (97% of 24866 models), with a model precision of 67Å. Model precision is the variability of models in this cluster and is computed as the average RMSD of the cluster models to the cluster centroid (Fig 3, Fig. S2, Methods). The model precision is lower than the resolution of the ODP tomogram (32 Å) (Al-Amoudi et al., 2011). This is mostly due to the fact that the integrative model localizes 55% more residues than the tomogram, the majority of which are on disordered and flexible regions. Other factors that contribute to low precision include low-resolution, sparse, and noisy input information. For example, the protein-protein binding data is on the domain level and not the residue level, we have fewer than ten protein-protein binding restraints, and the immuno-EM data has large error bars. All these sources of uncertainty in the input information are reflected in the model precision.

**Figure 3.**
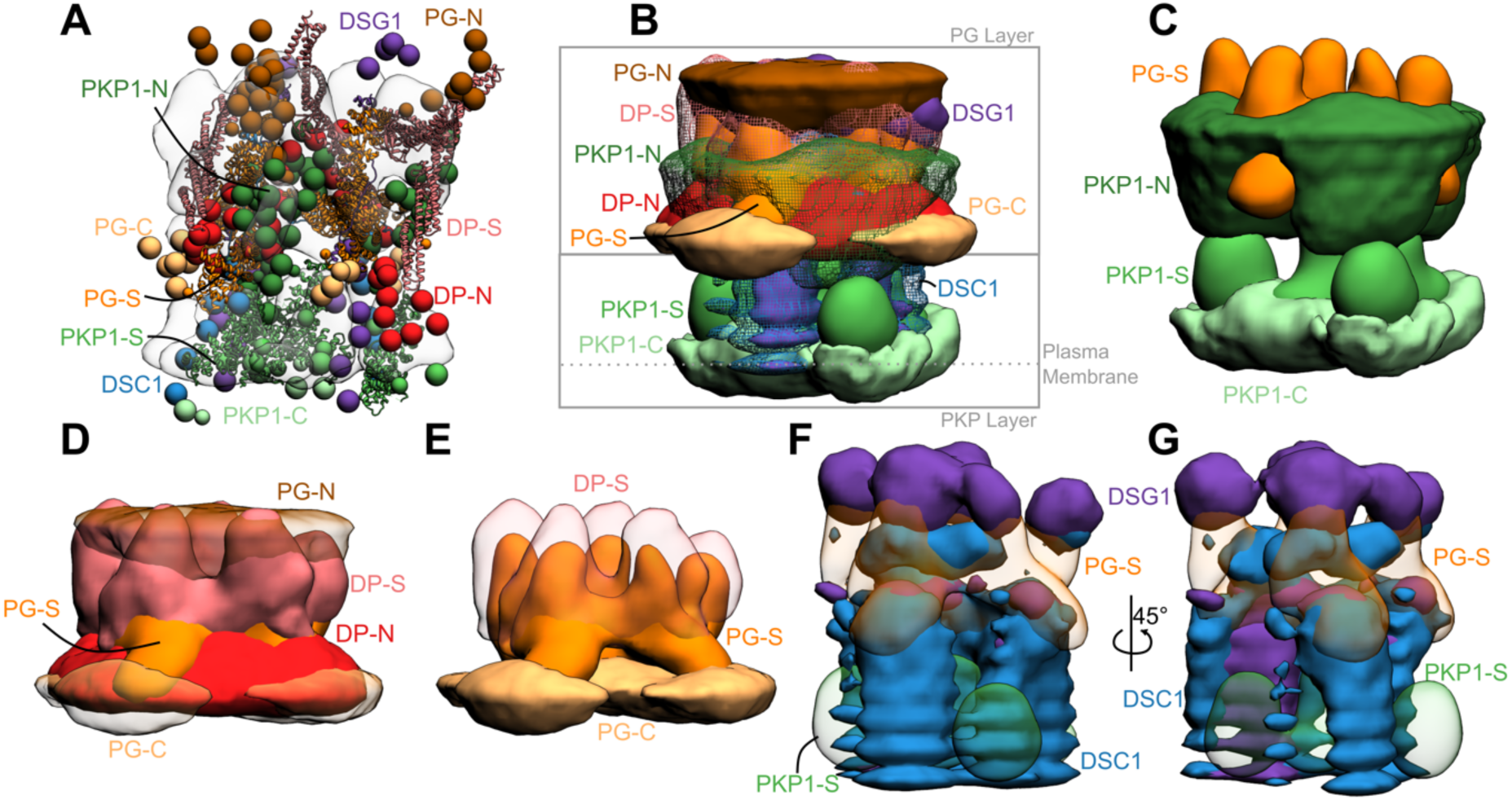
Integrative structure of the desmosomal ODP. **A)** The cluster center bead model for the major structural cluster with the cryo-tomogram (EMD-1703) superimposed in translucent gray. **B)** Localization densities of the major cluster. The densities are at a cutoff of approximately 15% for PKP1-C, PKP1-S, PG-S, DP-S, DSC1, DSG1 and around 30% for disordered termini regions (PKP1-N, PG-N, DP-N, PG-C). **C)** Localization densities for PKP1 layer (PG-S density is shown for reference). **D)** Localization densities for PG-layer. **E)** The densities for PG-S and DP-S with PG-C as a reference. **F-G)** Localization densities for the cadherins. Panel G is a rotated view of Panel F. See also Fig. S2-S4.

Nevertheless, these models fit well the input information used in modeling (Fig. S3, Table S2, Methods). They were further corroborated by their excellent agreement with information not used for modeling (Fig. S4, Table S3). The resulting integrative structures were visualized in two ways: a bead model representing the centroid of the major cluster (Fig. 3A), and a localization probability density map, representing the localization of protein domains by specifying the probability of a voxel (3D volume) being occupied by a domain in the set of structurally superposed cluster models (Fig. 3B-3G).

Overall, the desmosomal ODP resembles a densely packed cylinder with two layers, the PG layer on top of the PKP layer (Fig. 3A-3B). A striking feature of the ODP model is that the two layers are not distinct and well-separated. Rather, the desmosomal cadherins and PKP1 span both the layers. The N-terminus of PKP1 penetrates into the PG layer while the rest of it is in the PKP layer (Fig. 3B).

#### PKP layer

PKP1-C is the region of the ODP closest to the plasma membrane. This region has low precision in the integrative model as shown by the spread of the localization densities (Fig. 3B-3C). PKP1-S, the armadillo repeat domain of PKP1, is juxtaposed between PKP1-C and PKP1-N, at high precision (Fig. 3B-3C). This is consistent with PKP1-S localization in tomograms (Al-Amoudi et al., 2011). PKP1-N extends from PKP1-S in the PKP layer to the middle of the PG layer, forming interfaces with several proteins in the PG layer (see also Protein-protein interfaces) (Fig. 3B-3C). Its density is spread out, *i.e.*, it has a low precision, consistent with the idea that it is a disordered domain (Fig. 3B-3C) (Al-Amoudi et al., 2011).

#### PG layer

PKP1-N, DP-N, and PG-C form the approximate boundary between the PKP and PG layers (Fig. 3B-3D). The last two are approximately equidistant from the plasma membrane, consistent with previous immuno-EM studies (North et al., 1999)(Table S2B). PG-S and DP-S, the armadillo repeat and plakin domains of PG and DP respectively, seem to localize in approximately the same region and physically interact (see also Protein-protein interfaces) (Fig. 3B-3E). Previously, PG-S and DP-S were hypothesized to form a regular zigzag arrangement, with both domains approximately equi-distant to the plasma membrane (Al-Amoudi et al., 2011). In contrast, in our integrative structure, the centers of PG-S and DP-S are at slightly different distances from the membrane (Fig. 3D-3E). On average, PG-S is slightly closer to the plasma membrane and DP-S is slightly closer to the cytoplasmic end. Also, there is no regular orientation to either PG-S or DP-S, although based on the localization densities, these domains appear to prefer an orientation where their long axis is approximately perpendicular to the membrane (Fig. 3D-3E). The lack of regular orientations could be because these domains are flexible and dynamic. Alternatively, the orientation could be regular, but there is not enough data at present to suggest a regular orientation.

The cytoplasmic end of the desmosomal ODP is occupied by PG-N. The PG layer protein termini having unstructured domains, PG-N, DP-N, and PG-C, are localized at low precision (Fig. 3D).

#### Desmosomal cadherins

The desmosomal cadherins extend from the membrane end of the ODP, through the space in the PKP layer, towards the PG layer, interacting with PG, DP, as well as PKP1 (see also Protein-protein interfaces) (Fig. 3B, 3F-3G). DSG1 being longer, extends towards the cytoplasmic end of the PG layer, close to PG-N, where it is localized at low precision. Whereas, DSC1 extends until PG-S in the middle of the PG layer (Fig. 3B, 3F-3G).

### Protein-protein interfaces

To enable the discovery of protein-protein interfaces in the desmosomal ODP, we computed contact maps and predicted interfaces between protein pairs (Fig. 4, Fig. S5, Table S4, Methods). Our contact maps denote the percentage of models in the cluster in which the corresponding bead surfaces are within contact distance (10Å). The contact maps are consistent with the localization of PG and PKP1 in separate layers and with the structures of known ODP sub-complexes, *e.g.*, the PG-desmosomal cadherin complexes (Fig. S5, Table S4). Analysis of the set of top 2-5% contacts, which likely excludes contacts made randomly, enabled us to identify refine previously known interactions (Fig. 4, Fig. S5, Table S4, Methods). In general, the newly predicted interfaces are consistent with input biochemical binding information and refine the latter, providing higher-resolution information due to the integration of additional sources of information in the modeling. They form an extensive set of concrete hypotheses for future experiments (Fig. 4, Fig. S5, Table S4). Below, we discuss some of these novel interfaces in light of the role of desmosomal subunits in maintaining robust cell-cell adhesion, assembly of desmosomes, and desmosome-related diseases.

**Figure 4.**
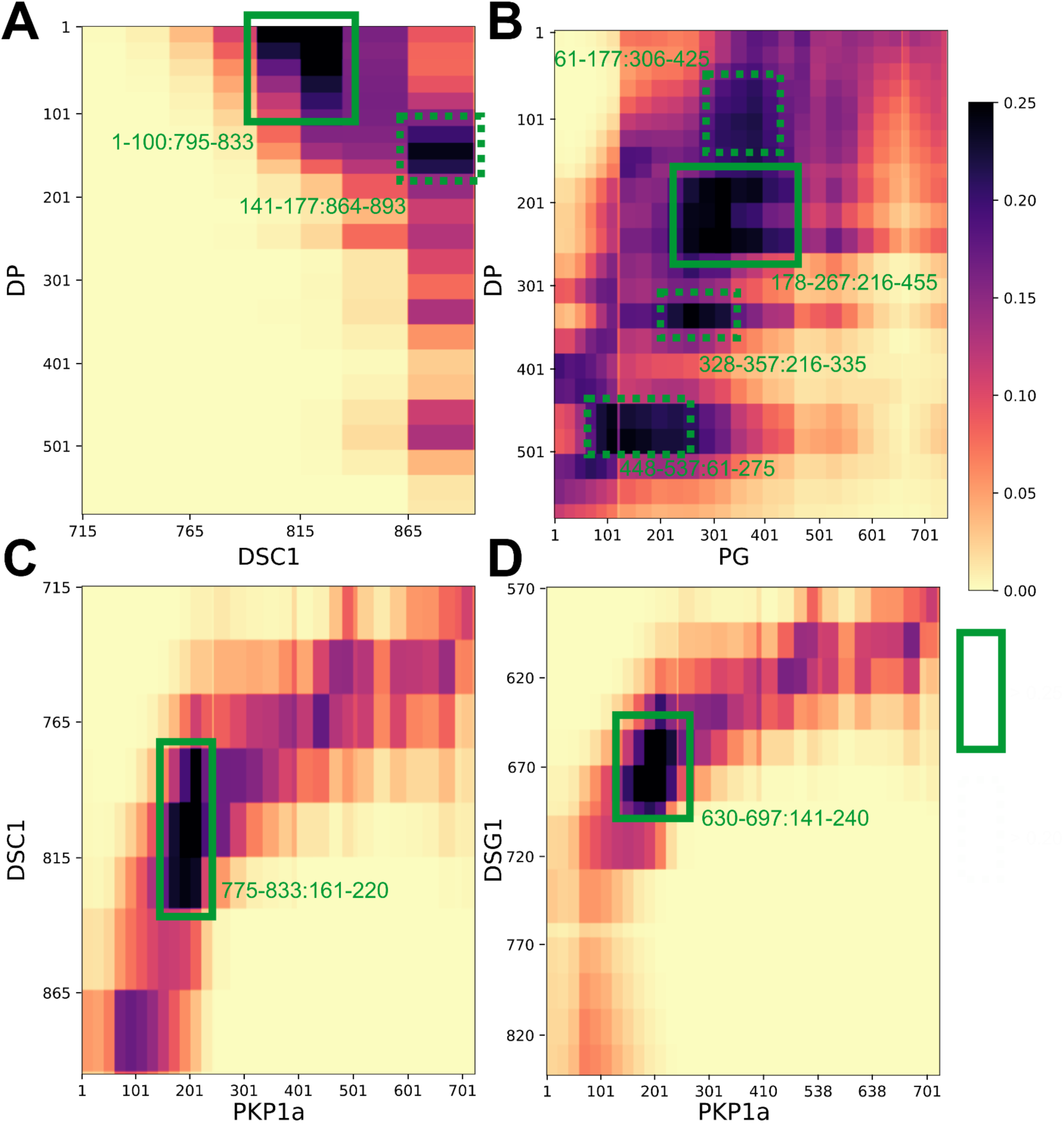
Novel ODP protein-protein interfaces. Protein-protein contact maps for DP-DSC1 (**A**), DP-PG (**B**), DSC1-PKP1 (**C**), and DSG1-PKP1 (**D**) pairs. Maps are colored by the proportion of the models in the major cluster where the corresponding bead surfaces are within contact distance (10 Å). Rectangles with solid green (broken green) lines outline novel contacts present in >25% (>20%) of the models. See also Fig. S5, Table S4.

### Insights into the molecular basis of desmosome-related diseases

Next, we hypothesized the structural basis for desmosomal defects in skin diseases and cancer by mapping disease-associated mutations on our integrative structure. These hypotheses would need to be verified experimentally in future studies. Specifically, we mapped known pathogenic missense mutations on desmosomal subunits that are associated with Naxos disease, Carvajal syndrome, or cancers (Fig. 5, Table S5, Methods). Both Naxos disease and Carvajal syndrome are characterized by abnormalities in epithelial tissue including palmoplantar keratoderma (thickened skin) and woolly hair (Boulé et al., 2012; Den Haan et al., 2009; Erken et al., 2011; Keller et al., 2012; Marino et al., 2017; McKoy et al., 2000; Pigors et al., 2015; Whittock et al., 2002).

**Figure 5.**
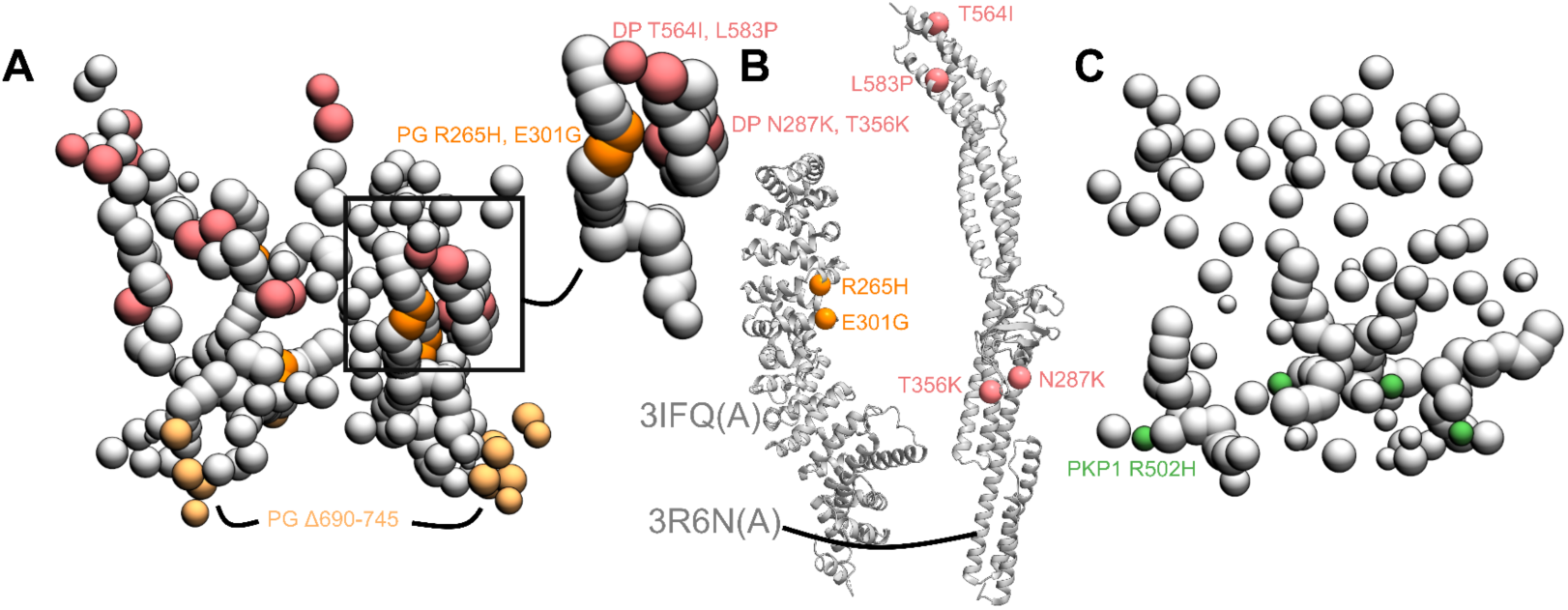
Disease-associated mutations mapped onto the integrative structure. **A)** Cluster center bead model showing mutations in PG and DP. Mutations in DP-S (pink), PG-S (orange) and PG-C (light orange) are colored as per Fig. 1. Remaining beads of DP and PG are shown in gray. Top right shows a zoomed-in version of a novel predicted PG-DP interface harboring disease mutations. **B)** PG-S and DP-S mutations mapped onto the corresponding structures 3IFQ(A) and 3R6N(A) (Choi et al., 2009; Choi & Weis, 2011) **C)** Bead model showing mutations in PKP1-S (green). See also Table S5.

#### PG mutations in Naxos Disease

The missense mutations PG R265H and PG E301G seen in Naxos disease are in the armadillo repeat domain of PG (Fig. 5A-5B, Table S5). These mutations are in the newly predicted PG-DP interface and known PG-DSG1 interface, and may result in disruption of these interfaces (Fig. 4, Fig. 5A-5B). Additionally, since they are in the armadillo domain, these mutations may also affect the folding and stability of this domain, and therefore desmosome assembly.

On the other hand, the truncation mutation PG Δ690-745 is in the disordered PG C-terminus (Fig. 5A). The latter is known to regulate the size of the desmosome; deletion of PG-C results in desmosomes that are larger than usual (Palka & Green, 1997). This truncation mutation may therefore affect desmosome assembly by altering the mechanism by which PG-C regulates desmosome size, *e.g.*, by modifying interactions with regulatory proteins.

#### DP mutations in Carvajal Syndrome and Skin Fragility/Woolly Hair (SF/WH) Syndrome

The DP missense mutations N287K (SF/WH syndrome) and T356K, T564I, and L583P (Carvajal Syndrome) are in the spectrin homology domain of DP (Fig. 5A-5B, Table S5) as well as the newly predicted PG-DP interface (Fig, 4, Fig. 5A, Table S4). These mutations may alter the integrity of the DP-PG interface as well as the folding and stability of the spectrin domain.

#### Cancers

The PKP1 R502H missense mutation is in the armadillo repeat domain of PKP1 and might affect the folding and stability of PKP1 in the ODP (Fig. 5C, Table S5). It is noteworthy that this residue is missing in the PDB structure of PKP1, PDB: 1XM9 (Choi & Weis, 2005)), indicating that it could be heterogeneous.

The other mutations associated with these diseases could not be readily rationalized by our structure (Table S5). In summary, three reasons can be identified for the pathogenicity of these mutations. They alter the folding and/or stability of ODP proteins, they disrupt protein-protein interfaces in the ODP, or they modify the binding properties of functionally important disordered protein domains in the ODP. All three types of mutations may disrupt the assembly and stability of the ODP, thereby affecting cell-cell adhesion. However, these mutations could also be pathogenic due to their effects on other functions such as cell signalling (Garrod & Chidgey, 2008).

Additionally, the structure of cardiac desmosomes is likely similar to that of the modeled epithelial desmosome. Therefore, our model could also be used to determine the structural basis of the numerous mutations related to cardiac diseases (*e.g.*, ARVC). However, we restricted the mutation analysis to epithelial diseases since our integrative structure is based on epithelial tissue isoforms (cardiac tissues consist of a slightly different set of isoforms (Delva et al., 2009; Green & Simpson, 2007)) (Fig. 1, Table S1).

### PKP-N penetrates to the PG layer

Our models indicate that PKP1-N penetrates to the PG layer and a conserved forty-residue segment in PKP1-N interacts with several ODP proteins. In our integrative structure, the N-terminus of PKP1 (PKP1-N) penetrates from the PKP layer to the PG layer and the two layers are not well-separated (Fig. 3A-3C, Fig. 6). In contrast, PG and PKP were seen in two distinct layers in cryo-electron tomograms (Al-Amoudi et al., 2011). The densities in these tomograms were likely contributed by regions of known structure (*e.g.*, PG-S and PKP1-S). PKP1-N, being disordered, is possibly flexible and heterogenous, leading to smoothing out of its densities upon averaging (Fig. S6). In integrative modeling *via* IMP, regions of unknown structure can be modeled alongside regions of known structure. By combining biochemical binding data along with structural (electron cryo-tomography) data, our approach allowed us to localize disordered domains like PKP1-N.

**Figure 6.**
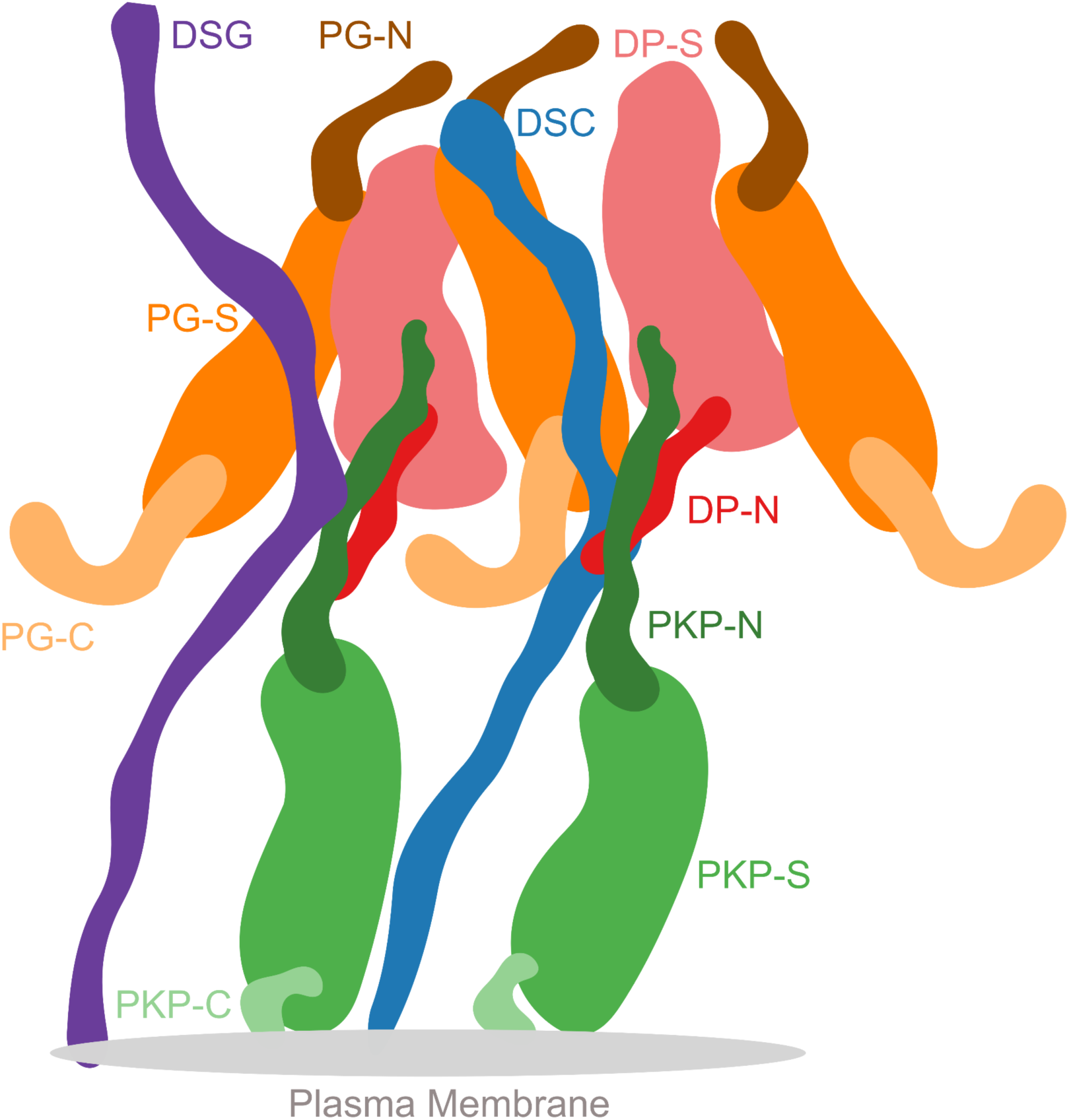
Schematic of the desmosome ODP. Schematic showing the salient features of the protein organization and protein-protein interfaces in the ODP. Wavy thick lines represent potentially disordered regions without known structure (DC, PKP-N, PG-N, DP-N, PKP-C, PG-C). Larger shapes represent regions with known structure (PG-S, DP-S, PKP-S). This is an artistic representation of Fig 3 which contains quantitative information from the model in terms of localization density maps.

In our structure, N-PKP mediates interactions with several ODP proteins (Fig. 4, Fig. 6), implying that PKP plays a more integral role in desmosome function and assembly. Specifically, PKP1^181-220^ interacts with DSC1^775-814^, and PKP1^161-220^ with DSG1^650-689^; notably, both desmosomal cadherins share binding sites on PKP1 (Fig. 4). Also, PKP1^141-180^ interacts with DP^1-60^ at a slightly lower, but still stringent, contact map cutoff (top 5% of all contacts, Fig. S5, Table S3). Interestingly, this forty residue stretch in PKP1-N, PKP1^161-200^, interacts with DP as well as the cadherins, and sequence analysis suggests that this sequence in PKP1-N is conserved (Fig. S6).

This is consistent with studies that show that PKP1 enhances recruitment of other desmosomal proteins, increasing desmosome size, and promoting desmosome assembly. For example, (Bornslaeger et al., 2001; Kowalczyk et al., 1999; Sobolik-Delmaire et al., 2006) showed that PKP1 clusters DP, (Hatzfeld et al., 2000) showed that PKP1 interacts with DP as well as desmosomal cadherins, and keratins, and (Tucker et al., 2014) showed that PKP1 interacts with DP and DSG3. PKP1 is also essential for clustering DSG1 and DSG3 (Fuchs et al., 2019). Two of the above studies mention that the amino tip of PKP1, PKP1^1-70^ (Hatzfeld et al., 2000) and PKP1^1-34^ (Sobolik-Delmaire et al., 2006), recruits DP. In our models, although PKP1^141-180^ (middle of PKP1-N), is the most probable PKP1-binding region (*i.e.,* highest confidence contact) for DP, the amino-tip of PKP1 is also proximal to DP and is among the top 5% of PKP1-DP contacts (Table S4). This region is also predicted to bind to DP based on Alphafold2-Multimer (See Comparison to Alphafold2-Multimer). In summary, our models indicate that PKP1-N, specifically the conserved region PKP1^161-200^, could be involved in the recruitment and/or subsequent stabilization of other ODP proteins.

Further, it is seen that PKP2 is highly mobile and exhibits fast turnover, whereas desmosomal cadherins, PG, and DP form a stable module from FRAP experiments (Fülle et al., 2021). This is in line with our findings: there are several contacts on the stable module, whereas the majority of PKP, *e.g.*, PKP-S did not appear to have contacts with other desmosomal proteins (Fig. 3, 4, Table S2, S4). This could imply that plakophilins play a signalling role in the desmosome while PG, DP, and cadherins form a structurally stable core.

### PG-C extends outwards from the ODP

In our integrative structure, the C-terminus of PG, PG^674-745^, extends outwards, suggesting that it can form lateral connections with other proteins (Fig. 3D, Fig. 6). It is known to play a role in regulating the size of the desmosome. Deletion of the PG C-terminus resulted in larger desmosomes due to lateral association (Palka & Green, 1997). Moreover, this deletion was also associated with Naxos disease and defects in tissue integrity, highlighting the importance of PG-C (McKoy et al., 2000).

The mechanism by which PG-C regulates the size of desmosomes remains to be elucidated. It is predicted to be intrinsically disordered (IDR) (Fig. S6). PG^683-687^ in this region is predicted to be a MoRF (molecular recognition feature), which is a motif in a disordered protein sequence that recognizes and binds to another protein (Disfani et al., 2012). The presence of the MoRF may allow PG-C to bind to itself, *i.e.,* PG-S, or to other proteins to enable regulation of desmosome size. In particular, the former mechanism, *i.e.*, IDR tails competitively binding to domains of the same protein to inhibit their function, is well-known for several enzymes and single-stranded DNA-binding proteins (Uversky, 2013). Finally, this region also contains a phosphosite (PG S730), suggesting that phosphorylation could potentially be another mechanism by which the desmosome size is regulated (Bian et al., 2014).

### Plakin domain of DP interacts with the armadillo repeats of PG

Our integrative structure identifies an interaction between the plakin domain of DP and the armadillo repeat domain of PG, DP^178-267^ and PG^276-335^ (Fig. 4). DP-S appears to encapsulate PG-S in the densities (Fig. 3E, Fig. 6). This interaction could provide a robust mechanism for desmosomes to anchor intermediate filaments (IF) and withstand mechanical stress. In fact, PG-DP binding is shown to be required for effective IF anchoring in desmosomes. PG knockout cells showed defective anchoring of IF (Acehan et al., 2008). Both the DP plakin domain and the PG arm domain are conserved across vertebrates, suggesting this interaction could also be conserved (Green et al., 2020; Smith & Fuchs, 1998). Further, a mutation in this region, PG E301G, was associated with Naxos disease and defects in epithelial tissue, further alluding to the importance of this interaction (Fig. 5, Table S5).

Moreover, this interaction could be important for desmosome assembly. In transient expression experiments in COS cells, PG was shown to be required for DP recruitment to cell borders (Kowalczyk et al., 1999). In our models, the DP binding region of PG overlaps with its cadherin-binding region, consistent with the fact that these three ODP proteins cluster together in desmosome assembly (Fig. 4) (Kowalczyk et al., 1999). Given their proximity, DP could also regulate the signaling functions of PG and PG-mediated crosstalk between desmosomes and adherens junction (Garrod & Chidgey, 2008).

### Localization and interactions of desmosomal cadherins

The desmosomal cadherins wind their way through the other proteins in the PG and PKP layers, making several interactions (Fig. 3B, F, G, Fig. 6). The cadherin spacing of 7nm from recent cryo-electron tomograms is consistent with our model (Sikora et al., 2020)(Fig. S4, Methods). The cadherins appear to be embedded in the thick of the other proteins, instead of circumnavigating the other proteins. This embedding in the midst of other ODP proteins provides a stronger anchoring for the cadherins and their extracellular domains in the cytoplasm. In turn, this feature could buffer the desmosomes from mechanical stress.

Notably, DSC and DSG different in their interactions with the other proteins. DSC1^795-833^ (the DSC1 region N-terminal to its PG binding site) interacts with DP^1-60^ (Fig. 4). Whereas, an interaction with DP is not seen for DSG1 (Fig. S5). This is consistent with the input information (Fig. 1, Table S2) (Smith & Fuchs, 1998). It is also consistent with experiments that showed that DSG requires PG to recruit DP, while DSC can recruit DP independently (Kowalczyk et al., 1999).

### Comparison to Alphafold2-Multimer

We also attempted to model sub-complexes of the ODP using the recent Al-based protein structure prediction method, Alphafold2-Multimer (Evans et al., 2021)(Methods Stage 4).

#### PG-DSC1 and PG-DSG1 complexes

AF2-multimer correctly reproduced the PG-DSC1 and PG-DSG1 complexes which were homology modeled based on PDB 3IFQ in this study (Fig. 1, Fig. 4, Fig. S7A-S7B, Fig. S8A-S8B). The template was likely part of the AF2 training set (all pre-2019 PDB structures).

#### PKP1-DSC1 and PKP1-DSG1 complexes

AF2-Multimer produced confident predictions for the PKP1-DSG1 and PKP1-DSC1 complexes (Fig. S7C-S7D, Fig. S8C-S8D). In these predictions, the PKP1 binding region for DSG1 and DSC1 is approximately similar to the PG binding region for cadherins, which is a reasonable prediction based on structural similarity, since PKP1 has an armadillo domain like PG (Fig, 4, Fig. S7C-S7D, Fig. S8C-S8D).

The DSC1 region binding to PKP1 is distinct from that binding to PG and is consistent with the region of DSC1 located in the PKP layer in our model. However, the DSG1 region that binds PKP1 in the Alphafold2 structure overlaps with its PG-binding region in our model. Additionally, our contact predictions from integrative modeling identify interfaces between the disordered PKP1-N and desmosomal cadherins, which are not captured in AF2-Multimer.

#### PKP1-DP and PG-DP complexes

Interestingly, AF2 predicted an interface between a part of the disordered N-terminus of PKP1 (approximately PKP1^20-51^) and DP (Fig. S7E, Fig. S8E), predicting a potential disordered-to-ordered transition on binding for PKP1. The predicted interface overlaps with our contact map predictions of interfaces between DP and PKP1 (Fig. 4, Fig. S5, Table S4) and is also consistent with studies that show that the tip of PKP1-N binds to DP (Hatzfeld et al., 2000; Sobolik-Delmaire et al., 2006). However, there was no predicted interface between PG and DP.

AF2-multimer presumably predicts the structure of a complex if it is similar to known complexes, or if it involves disordered regions that become ordered upon binding to a partner. However, the predicted interface information is incomplete. For example, no interface was detected for the PG-DP complex. AF2-multimer predicts a single model or a small number of candidate models, while our integrative modeling method predicts a larger probability-weighted ensemble of models consistent with input information. Furthermore, it is a deep-learning method based on general patterns in existing protein structures. It does not account for information that is specific to a system, such as the membrane topology, layered arrangement of proteins or the oligomeric states of proteins. Lastly, these are only dimeric predictions, and the error in AF2-Multimer predictions would get amplified for larger multimeric assemblies, such as the full desmosomal ODP, leading to a potentially inaccurate prediction. In summary, tools like AF2-Multimer are not currently sufficient to model the complete desmosome at high-resolution, presumably due to low sequence similarity to existing structures and the presence of disordered regions.

Here, we obtained an integrative structure of the desmosomal ODP starting primarily from the structural interpretation of an electron cryo-tomogram of the ODP, and combining X-ray crystal structures, distances from an immuno-EM study, interacting protein domains from biochemical assays, bioinformatics sequence-based predictions of transmembrane and disordered regions, homology modeling, and stereochemistry information. High-resolution structural data, *e.g.*, higher-resolution cryo-EM maps would improve the structural characterization of the desmosome and our knowledge of the mechanistic details of cell-cell adhesion. Structural characterization of the desmosome interactome, for example, desmosome-associated adaptor proteins is another avenue for future work (Badu-Nkansah & Lechler, 2020).

## Methods

Integrative structure determination of the desmosomal ODP proceeded through four stages (Alber et al., 2007; Rout & Sali, 2019) (Fig. 1-2). Our modeling procedure used the Python Modeling Interface of the Integrative Modeling Platform (IMP 2.17.0; https://integrativemodeling.org), an open-source library for modeling macromolecular complexes (Russel et al., 2012), and is primarily based on previously described protocols (Arvindekar et al., 2022; Saltzberg et al., 2021; Viswanath, Chemmama, et al., 2017). Python libraries scipy (Virtanen et al., 2020) and matplotlib (Hunter, 2007) were used for analysis, GNU Parallel (Tange, Ole, 2020) was used for parallelization, UCSF Chimera v1.15 (Pettersen et al., 2004) and UCSF ChimeraX v1.5 (Pettersen et al., 2021) were used for visualization. Input data, scripts, and results are publicly available at https://github.com/isblab/desmosome and ZENODO. Integrative structures will be deposited in the PDB-DEV (https://pdb-dev.wwpdb.org) .

### Stage 1: Gathering data

#### Isoforms

The ODP comprises of PG (plakoglobin), PKP (plakophilin), DP (desmoplakin), and Desmosomal Cadherins (DC of two types, Desmoglein, DSG, and Desmocollin, DSC). Desmosomes from different tissues vary in the isoforms of these constituent proteins (Garrod & Chidgey, 2008; Green & Simpson, 2007). Here, we modeled the desmosomal ODPs corresponding to the stratified epithelium and containing PKP1 (Fig. 1A, Table S1). For ODPs from two other tissues that we modeled (stratified epithelium containing PKP3, DSC1, DSG1, PG and DP and basal epithelium containing PKP3, DSC2, DSG3, PG, and DP) the results were similar at the resolution of the input information (Fig. 3). Epithelial desmosomes were chosen for modeling as there was more information (*e.g.*, from protein-protein binding experiments) on epithelial isoforms than desmosomes in Heart tissue. The extracellular regions of the Desmosomal Cadherins were not modeled, based on sequence annotations in Uniprot (see also (Choi et al., 2009)). Further, we do not model DSG1^843-1049^ and DP^585-2871^ as they are known to be outside the ODP (Al-Amoudi et al., 2011; Garrod & Chidgey, 2008; Nilles et al., 1991)(Table S1).

#### Stoichiometry and number of copies

The stoichiometry of the desmosomal proteins was based on previous studies using modeling and density analysis on cryo-electron microscopy data (Al-Amoudi et al., 2011)(See Stage 2).

#### Atomic structures

The plakin domain of DP and armadillo domains of PG and PKP1 were modeled by their X-ray structures (PDB: 1XM9 (PKP) (Choi & Weis, 2005), 3R6N (DP) (Choi & Weis, 2011), 3IFQ (PG-DC) (Choi et al., 2009), while the PG-DSC and PG-DSG complexes were obtained by homology modeling using MODELLER (Šali & Blundell, 1993) and HHPRED (Gabler et al., 2020) for sequence alignment (Fig. 1C, Table S1).

#### Cryo-electron tomogram

We used a 32 Å cryo-electron tomogram (EMD-1703, denoised mask without symmetrization) of the ODP (Al-Amoudi et al., 2011). The map was segmented using UCSF Chimera Segger (Pintilie et al., 2010) and the densities corresponding to the PKP and PG layers were used for modeling. (Fig. 1C).

#### Immuno-EM

The distance of the N and C termini of the desmosomal proteins from the plasma membrane was informed by immuno-electron microscopy gold-staining experiments (Fig. 1B, Table S2) (North et al., 1999). Using Clustal-Omega (Sievers et al., 2011), the alignment between Bovine/Xenopus PG and DP (used in the experiments) and the Human PG and DP (used in modeling) is almost 1-to-1, and therefore, the residue ranges for the antibody-binding regions are taken to be the same. Values for PKP1 were used for PKP3 after alignment.

#### Protein-protein binding assays

The relative distance between ODP protein domains was informed by biochemical data from multiple biochemical studies, including yeast-2-hybrid (Bonné et al., 2003; Hatzfeld et al., 2000; Kowalczyk et al., 1999), co-immunoprecipitation (Bonné et al., 2003; Kowalczyk et al., 1999), *in-vitro* overlay assays (Smith & Fuchs, 1998), and *in-vivo* co-localisation assays (Bonné et al., 2003; Bornslaeger et al., 2001; Kowalczyk et al., 1999) (Fig.1B, Table S2-S3). Due to experimental issues, the information pertaining to DSC3a binding is not usable from (Bonné et al., 2003) and we therefore use the corresponding information from DSC3b binding.

We note that we have not used other desmosome data that is not directly informative for the integrative structural modeling of the core stratified epithelial ODP. This includes data on desmosome-interacting proteins, data on isoforms of ODP proteins not dominant in the modeled tissue, data on the role of desmosome in signaling and regulation, and data that is too low-resolution for our modeling (for example, on the protein level instead of the domain or residue level).

### Stage 2: Representing the system and translating data into spatial restraints

#### Stoichiometry and number of copies, PG layer and the Desmosomal Cadherins

The stoichiometry of the desmosome ODP was 1:1:1:1 for DP:PG:PKP:DC based on previous studies (Al-Amoudi et al., 2011). The number of copies of each protein was based on fitting an equal number of PG and DP molecules to the PG layer of the cryo-electron tomogram. However, the number of PG and DP proteins that correspond to the tomogram was unknown and computed to be four each by fitting different numbers of PG and DP molecules to the PG layer density in independent modeling runs (Supplementary Section 1.1). We model 4 DC molecules, two each of DSC1 and DSG1.

#### Stoichiometry and number of copies, PKP layer

The PKP layer has seven distinct densities. These correspond well (average EM cross-correlation around mean in UCSF Chimera = 0.91) to the structured ARM repeats of seven PKP molecules (Al-Amoudi et al., 2011)(Fig. S1 inset). To keep a 1:1:1:1 stoichiometry for PG:DP:PKP:DC, we selected four of these seven PKP molecules to represent in full; the central PKP and three symmetrically surrounding PKPs (Fig. S1 inset).

We also represented the remaining three PKP molecules (“non-interacting” PKPs) by their structured ARM repeats alone. These PKPs participate only to satisfy the cryo-electron tomogram and to exclude other proteins from these locations in space. The locations and orientations of each of these PKPs were fitted based on cross-correlation to the PKP densities in the tomogram; subsequently they were fixed during sampling.

#### Multi-scale coarse-grained bead representation

The rationale for choosing a coarse-grained representation is based on the following requirements (Viswanath & Sali, 2019). A representation must enable efficient and exhaustive sampling of models, its resolution must be commensurate with the quantity and resolution of input information, and the resulting models should facilitate downstream biological analysis. In the current case, we have low resolution, sparse, noisy sparse input information, therefore, a higher-resolution representation, for example, 1-residue per bead representation, would not be justified. Moreover, sampling with this higher resolution representation would be infeasible in days on modern supercomputers for a complex as large as the ODP; on the other hand, sampling in a shorter time would not be exhaustive.

In light of these considerations, we use the following coarse-grained representation of the proteins where a set of contiguous amino acids in a protein is represented by a spherical bead (Fig. 1A, Table S1). Domains with known atomic structures were represented by 30-residue beads to maximize computational efficiency and modeled as rigid bodies where the relative configuration between the beads is fixed during sampling. Notably, this coarse-graining of domains with known atomic structures is performed mainly for sampling efficiency and does not result in any loss of existing atomic structural information, as one can map these structures readily on to the rigid bodies in our model (Fig. 3). In contrast, domains without known structure were coarse-grained at 20 residues per bead and modeled as flexible strings of beads which can move relative to one another.

Next, we encoded the information gathered in stage 1 into spatial restraints that constitute a scoring function which allows scoring each model in proportion to its posterior probability. This score allows sampling high-probability models that best satisfy the data.

#### EM restraints

A Bayesian EM restraint was used to incorporate the information from the cryo-electron tomogram (Bonomi et al., 2019). PKP-S, the structured region of PKP, was restrained by the PKP-layer density; PG and DP molecules were restrained by the PG layer density. The EM restraint was not applied to regions such as PKP-N, PKP-C, and the desmosomal cadherins as they are either disordered and/or extended and therefore considered to be averaged out or contribute negligibly to the density in the tomogram (Al-Amoudi et al., 2011). The part of DC complexed with PG was included in this restraint.

#### Immuno-EM restraints

The distances of ODP protein termini to the plasma membrane were restrained by a Gaussian restraint with the mean and standard deviation equal to the mean distance and standard error measured in immuno-EM gold-staining experiments (North et al., 1999). The set of restrained beads for each protein terminus corresponded to the antibody-binding region in the experiments. The standard error of mean accounts for the variance in the distance measurements arising, for example, from random antibody orientations. The restraint score was based on the bead in the terminus that was closest to the mean distance obtained from the experiment, for each protein copy. Desmosomal Cadherins were not restrained by immuno-EM since they form a complex with PG, which is restrained by immuno-EM data. The complexed region is more specific than the antibody-binding region for DSG1. Further, immuno-EM measurements were not available for specific DSC isoforms.

#### Binding restraints

The distances between interacting protein domains were restrained by a harmonic upper bound on the minimum distance among the pairs of beads representing the two interacting domains, (the score is zero for distances less than or equal to 0, and quadratically rises above zero). For two interacting proteins A and B, ambiguity, *i.e.,* multiple copies of a protein, was factored in by adding multiple such distance restraints. For each copy of protein A, the minimum distance among all pairs of beads across all copies of B was restrained. Similarly, for each copy of protein B, the minimum distance among all pairs of beads across all copies of A was restrained. This formulation allows a protein copy to find a binding partner from any of the available copies of the other protein, potentially allowing multiple protein A copies to bind to the same protein B copy.

Different experiments provide different levels of evidence as to whether their results can be extended to *in-vivo* conditions and whether the results preclude indirect binding via an intermediary protein. Restraints were therefore weighed in the order Overlay Assays = Co-Immunoprecipitation > Yeast-2-Hybrid. However, the results we obtain are fairly robust to this weighting scheme and all the experimental data are individually satisfied in the final set of models (Fig. S3). If multiple experiments provided data on the binding of two proteins, the highest-resolution data (*i.e.,* more specific binding site) was chosen.

#### Cylindrical restraints

To keep the modeled proteins close to the tomogram, beads were restrained to lie within a cylinder of radius 150 Å that encloses the map. The restraint was implemented using a harmonic upper bound on the distance of each bead from the cylinder surface.

#### Excluded volume restraints

The excluded volume restraints were applied to each bead to represent the steric hindrance of the protein residues that disallow other residues to come in physical proximity. The bead radius was calculated assuming standard protein density (Alber et al., 2007), with beads penalized based on the extent of their overlap.

#### Sequence connectivity restraints

We applied sequence connectivity restraints on the distance between consecutive beads in a protein molecule. The restraint was encoded as a harmonic upper bound score that penalizes beads that are greater than a threshold distance apart. The threshold distance is different for each protein and the calculation is inspired by models from statistical physics (Teraoka, 2002)(Supplementary Section 1.2). As a summary, we predict what proportion of each protein’s predicted secondary structure is disordered using PSIPRED (Buchan & Jones, 2019), and compute the threshold based on this proportion, the known radii of gyration for disordered regions, and bead radii for globular proteins estimated from their density (Alber et al., 2007). For regions with known structures, the inter-bead distances were fixed during sampling and their contribution to the restraint score was fixed across models.

### Stage 3: Structural sampling to produce an ensemble of structures that satisfies the restraints

We employed Gibbs sampling Replica Exchange Monte Carlo sampling (Arvindekar et al., 2022; Saltzberg et al., 2021; Viswanath, Chemmama, et al., 2017). The positions of the rigid bodies and flexible beads were sampled as in previous protocols, with a few customizations.

First, we implemented an Anchoring Constraint wherein the membrane-proximal beads of the desmosomal cadherins were initialized adjacent to the membrane and were constrained to move only along the membrane plane during sampling (Fig. 1B).

Second, a custom random initialization was used for the PG layer. The PG and DP rigid bodies and beads were randomized within a bounding box that tightly enclosed the PG layer density. The orientation of the long axis of the structured region of PG and DP molecules with respect to the membrane determines the polarity of each PG/DP molecule (N-to-C along the normal to the membrane). After the random initialization, if it was opposite of the polarity observed from immuno-EM (North et al., 1999), this polarity was corrected by flipping the structured region along a random axis in the plane of the membrane by 180 degrees; in effect, reversing the polarity along the normal to the membrane while keeping its orientation random. For example, if a PG molecule was initialized with its N-terminus closer to the membrane than its C-terminus, its orientation would be flipped. This is because, owing to the high protein density of the PG layer, molecules with the incorrect polarity might not have the freedom to flip polarity during sampling.

Finally, a custom random initialization was used for the PKP layer. Each PKP was initialized around one of the molecule-wise PKP densities with a random orientation.

The Monte Carlo moves included random translations of individual beads in the flexible segments, random translations and rotations of rigid bodies and super-rigid bodies, *i.e.*, groups of rigid bodies and beads of the same protein or complex. The size of these moves and the replica exchange temperature for the replicas was optimized using StOP (Pasani & Viswanath, 2021). A model was saved every 10 Gibbs sampling steps, each consisting of a cycle of Monte Carlo steps that proposed a move for every bead and rigid body once. We sampled a total of 180 million integrative models, from 50 independent runs.

### Stage 4: Analyzing and validating the ensemble of structures

The sampled models were analyzed to assess sampling exhaustiveness and estimate the precision of the structure, its consistency with input data and consistency with data not used in modeling. We based our analysis on the protocols published earlier (Arvindekar et al., 2022; Saltzberg et al., 2021; Viswanath, Chemmama, et al., 2017; Webb et al., 2018).

#### Filtering the models into a good-scoring set

To make analysis computationally tractable and to select models that have a good score, *i.e.* higher probability, we first selected the models to create a good-scoring set which involved the following steps. Models were first filtered based on score equilibration and auto-correlation decay along the MCMC runs (Supplementary Section 2.1). Filtered models were clustered based on their restraint scores using HDBSCAN (McInnes et al., 2017), resulting in a single cluster of 37145 models (Saltzberg et al., 2021). Subsequently, these models were filtered to choose models for which each restraint score as well as the total score is better than the corresponding mean plus 1.46 standard deviations, leading to a good-scoring set of 24866 models for the next stage of analysis (Arvindekar et al., 2022).

#### Clustering, Precision, and Localization Densities

We next assessed if the sampling was exhaustive by previously established protocols which randomly divide the models into two independent sets and assess *via* statistical tests whether the two sets had similar scores and structures (Arvindekar et al., 2022; Saltzberg et al., 2021; Viswanath, Chemmama, et al., 2017)(Fig S2). We perform structural clustering of the models to find the minimum clustering threshold for which the sampling is exhaustive (sampling precision) as well as the mean RMSD between a cluster model and its cluster centroid (model precision) (Fig. S2). The bead-wise RMSD calculation in the protocol was extended to consider ambiguity, *i.e.*, multiple protein copies, and the resulting code was parallelized. The RMSD between two models is the minimum RMSD among all combinations of protein copy pairings between the models. For example, two models containing four copies of PG have 4! possible bipartite pairings of PG copies among them for which the RMSD needs to be computed. The consideration of ambiguity was applied to all proteins except PKP. Each PKP copy was initialized to the same molecule-wise EM density in every simulation and usually remained close to it throughout the simulation. PKP copies could be considered as non-interchangeable because of the presence of fixed, non-interacting PKPs in their midst. The latter also precludes the need of alignment to a common frame of reference during RMSD calculations.

The result of integrative modeling was a single major cluster corresponding to 24016 (96.6% of 24866) models. The model precision, which quantifies the variability of models in the cluster, and is defined as the average RMSD of a cluster model from the cluster centroid, was 67Å. The cluster is visualized *via* localization probability density maps, which specify the probability of a volume element being occupied by beads of a given domain in the set of superposed models from the cluster (Fig 3).

#### Fit to input information

To calculate the fit to data from protein-protein binding assays, we calculated the minimum distance among all bipartite pairs of beads representing all copies of interacting domains for each model in the cluster and visualized the distribution (Fig. S3A, Fig. S4, Table S2A, Table S3A).

To calculate the fit to immuno-EM data (North et al., 1999) for each restrained protein terminus, we calculated the difference between the model-predicted distance of the terminus to the plasma membrane and the corresponding mean distance from experiment. The model-predicted distance for a terminus was equal to the distance of the terminus bead closest to the experimental mean. The distribution of the difference for each copy of a protein for each model in the cluster was visualized (Fig S3B).

To calculate the fit to the tomogram, we computed cross-correlation between the localization probability densities for the cluster and the segmented tomogram for the PG and PKP layers separately (Supplementary Section 2.2, Fig S3C). The model is consistent with all input information.

#### Fit to information not used in modeling

The fit to data from protein-protein binding assays was calculated similarly as above. With the exception of two experiments, the information not used in modeling was in complete agreement with our model (Fig. S4, Table S3B). The two exceptions correspond to data that mentions that the first few cytoplasmic residues of DSC1, in the PKP layer in our model, bind to proteins in the PG layer.

The fit to data from dSTORM super-resolution imaging was calculated as follows. As stated in (Stahley et al., 2016), the distances of modeled domains from the plasma membrane were obtained by starting from the dSTORM plaque-to-plaque measurements, subtracting the width of the intercellular space (∼34 nm, subtracting two times the plasma membrane thickness (∼4-6 nm), and dividing by two (Table S3B). These distances were compared to the distribution from our models which were already computed for comparing to the immuno-EM data as above (Fig. S3B, Table S3B). The dSTORM distribution of PG-N is consistent with (North et al., 1999) as well as our model (Fig. S3B, Table S3B). However, DP is localized further away from the membrane than it is in our model and in (North et al., 1999), a fact that Stahley and co-workers also comment on. This could be partly due to the uncertainty in dSTORM-based measurement that arises from the localization precision of Alexa Fluor 647, and the primary and secondary antibody labels.

To calculate the fit to data from electron cryo-tomography, we obtained the cadherin spacing from tomograms, *i.e.,* distance between DSG2 and DSC2 reported as 7 nm for the W-shape arrangement of cadherins and compared to the distribution from our model (Sikora et al., 2020)(Table S3B, Fig S4). The distribution of the minimum distance between the DSG1 and DSC1 membrane-anchored beads was plotted for models in the cluster, as a proxy for the distance between adjacent cadherins at the plasma membrane. The models are consistent with the spacing from these newer tomograms.

The models are also consistent by construction with the data from acyl biotin exchange assays from (Roberts et al., 2016) which state that DSG1 residues 571, 573 are membrane-proximal and palmitoylated (Table S3B, Fig. 1). The bead corresponding to residues 570-589 is membrane anchored in our model.

#### Contact Maps

A contact between beads is defined as a surface-to-surface distance of 10 Å or lower. For each protein pair, we obtained the proportion of models in the cluster that have at least one contact for each bead pair across all copies of the two proteins. To filter out the significant contacts from those that might occur by chance, we identified significant contacts as those present in at least 20-25% of the models. A 25% cutoff corresponds to approximately the top ≤2% of all possible contacts for each protein-pair while a 20% cutoff corresponds to ∼8% of all possible contacts for PG-DP and ≤5% for the rest of the protein pairs.

#### Mapping disease mutations

We considered two kinds of mutations to map to the integrative structure. First, disease mutations associated with defects in epithelial tissue that could be mapped to ODP protein domains and/or residues were obtained by a literature search and using databases such as OMIM and Uniprot (*Online Mendelian Inheritance in Man*, 2023; The UniProt Consortium et al., 2023). These mutations corresponded to those seen in Naxos disease (ARVC with palmoplantar keratoderma and woolly hair) and Carvajal syndrome (Left ventricular cardiomyopathy with palmoplantar keratoderma and woolly hair) (Boulé et al., 2012; Den Haan et al., 2009; Erken et al., 2011; Keller et al., 2012; Marino et al., 2017; McKoy et al., 2000; Pigors et al., 2015; Whittock et al., 2002). Second, cancer-associated somatic, missense, confirmed pathogenic mutations on ODP proteins that occurred in five or more samples were extracted from the COSMIC database (Tate et al., 2019).

We did not consider mutations involved in cardiac disease as we model an epithelial ODP. In general, we refrained from mapping mutations across isoforms. We also did not consider mutations that could not be mapped to the protein domains, although a large number of these are known, for example, pathological differential expression of proteins.

#### Comparison to Alphafold Multimer

We ran Alphafold2-Multimer (Evans et al., 2021) for pairs of proteins: PG-DP, PKP-DC, PG-DC and PKP-DP. For each pair, we chose the best ranked prediction, based on the PTM + IPTM score, to discover confidently predicted interfaces between the proteins. Confidently predicted interfaces were identified as residue pairs, with one residue from each protein, in which each residue was confidently predicted (pLDDT > 70), the residue pair had an accurate relative prediction (PAE < 5), and the pair was at an interface (Cα-Cα distance < 10Å).

## Data availability

Files containing input data, scripts, and results are at https://github.com/isblab/desmosome and at https://zenodo.org/doi/10.5281/zenodo.8035862. Integrative structures will be deposited in the PDB-Dev (https://pdb-dev.wwpdb.org).

## Acknowledgements

We thank lab members Shreyas Arvindekar, Kartik Majila, and Muskaan Jindal for comments on early versions of the draft. We thank Andrew Kowalczyk for helpful comments on the manuscript. We thank Aditi Pathak for help with Alphafold-multimer. We also thank Karan Gandhi, Sarika Tilwani, and Sorab Dalal of ACTREC, India and Swadhin Jana of NCBS for their help. Molecular graphics images were produced using the UCSF Chimera and UCSF ChimeraX packages from the Resource for Biocomputing, Visualization, and Informatics at the University of California, San Francisco (supported by NIH P41 RR001081, NIH R01-GM129325, and National Institute of Allergy and Infectious Diseases).

