## Supplementary Material for "The molecular architecture of the desmosomal outer dense plaque by integrative structural modeling"

### Section 1: Modeling

##### Section: 1.1 Stoichiometry Runs

The number of molecules that could be accommodated in the PG layer was unclear since it was not possible to unambiguously dock the molecules manually in this region. A higher number would result in an overcrowded model or push molecules outside the tomogram space and be spatially unconstrained by the tomogram, while a smaller number would not be enough to explain the whole EM map. To determine this, we ran independent modeling runs that only included the PG and DP molecules and the PG layer EM density. We ran six simulations, each with an equal number of PG and DP ranging from 2 to 7. For each of these runs, the representation and sampling followed the IMP modeling protocol (Saltzberg et al., 2021; Viswanath et al., 2017). The restraints applied included the EM restraint, the immuno-EM restraint, the excluded volume restraint, connectivity restraint, and the cylinder restraint to ensure that the molecules are not too far away from the tomogram. 3 million models were simulated per run. For each run, after filtering the models sampled before equilibration, the top 10% models were determined and the cross-correlation coefficient of these models to the cryo-electron tomogram was computed (Bonomi et al., 2019) (Fig. S1). The number of PG/DP molecules was taken to be four. It was only slightly lower in average cross-correlation than the value for three PG/DP molecules. Selecting four PG/DP copies allowed selecting four PKPs in the PKP layer including the central PKP without introducing any asymmetries in the selection. It also allowed us to maintain 1:1:1:1 stoichiometry for PG:DP:PKP:DC with equal numbers of DSC and DSG (2 each). Finally, it was consistent with previous studies which showed that no more than four PG and DP molecules each could fit in the PG layer (Al-Amoudi et al., 2011).

#### Section: 1.2 Restraints

##### Distance threshold for sequence connectivity restraint

To set up the connectivity restraint, we need to scale the inter-bead distance to allow the more disordered N/C termini as well as the DC proteins to span a greater end-to-end distance compared to the globular protein domains. For each protein domain with at least a partial disorder (for example, PKP-N, DP-N, etc), we first find the radius of gyration of this fragment assuming the fragment to be completely disordered, using $R_g = 1.92N^{0.6}$(Kohn et al., 2004) where *N* is the number of residues. We model the fragment as a chain of monomers composed of *n* statistically independent segments each of length *a*. We assume *n* to be approximately equal to the number of beads in our representation of the fragment as two adjacent beads are free to be in any relative orientation without any other restraints; *a* is then the inter-bead distance. We then use a relation between the RMS Distance between the two ends of the chain

($R_F = an^{0.6}$) and the radius of gyration ($R_G$) to estimate *a* for our fragment: ${R_G}^2/{R_F}^2 = 25/176$ (Teraoka, 2002). Another estimate for *a* is calculated internally in IMP and comes from the assumption that the fragment is globular (Alber et al., 2007). The final scaling depends on the weighted sum of these two estimates, the weights corresponding to the portion of the fragment predicted to be disordered by PSIPRED (Buchan & Jones, 2019). Given an estimate of *a*, we can calculate the surface-to-surface distance for adjacent beads to create an harmonic upper bound restraint such that the beads are only penalized when they are farther than this distance apart. We use the maximum end-to-end distance (*an*) and find the bead surface-to-surface distance needed to achieve this end-to-end distance. This is approximated by the following relation where *r* is the typical radius of a bead in our model: $d = (an - 2r)/n - 2r$. The calculated scale matches the scale calculated using a more accurate measure for ${R_G}^2/{R_F}^2 \approx 0.95/6$ given by renormalization theory (Teraoka, 2002) up to rounding. However, the scale is only a heuristic parameter and the results obtained are relatively robust to its exact value.

#### Section 2: Analysis

##### Section: 2.1 Filtering based on Autocorrelation

In order to filter a computationally feasible subset of models from the large set of sampled models, we first remove the initial few models based on  statistical testing (Chodera, 2016; Saltzberg et al., 2021), to consider only the models after equilibration assumed to be in the stationary distribution. Next, we only take every 20th model in the MCMC sampling run (PMI analysis parameter nskip=2, writing every 10th frame to disk). To identify an appropriate number of models to skip, we ran eight independent single-replica runs with all the restraints. We analyzed the spatial autocorrelation of the XYZ coordinates of each bead along the sampling trajectory. We chose as our cutoff the smallest number of sampling steps after which the autocorrelation of all the beads had fallen to at most 85-90%. This allows us to remove the highly correlated models to obtain an independent set of models to analyze downstream.

##### Section: 2.2 Cross-Correlation of Localization Densities and cryo-electron tomograms

We first computed the predicted localization density by combining the densities from all modeled proteins for the major cluster separately for the PKP layer (PKP-S) and PG-Layer (PG-N,S,C, DP-N,S). We then calculated the cross-correlation between the predicted density and the reference cryo-electron tomogram by calculating the Pearson correlation between the voxel-wise values in the two maps. This is calculated at all grid points at a voxel spacing of 5Å spread over the volume enclosing both the predicted localization density and the cryo-TM map. The values of the maps at these grid points were found by interpolation (RegularGridInterpolator in scipy (Virtanen et al., 2020)). This is similar to calculating Correlation around mean in UCSF Chimera (fitmap) except that the Chimera calculation only involves the non-zero grid points of the reference map, causing the correlation value to change depending on the order of the two maps.

#### Supplementary Figures

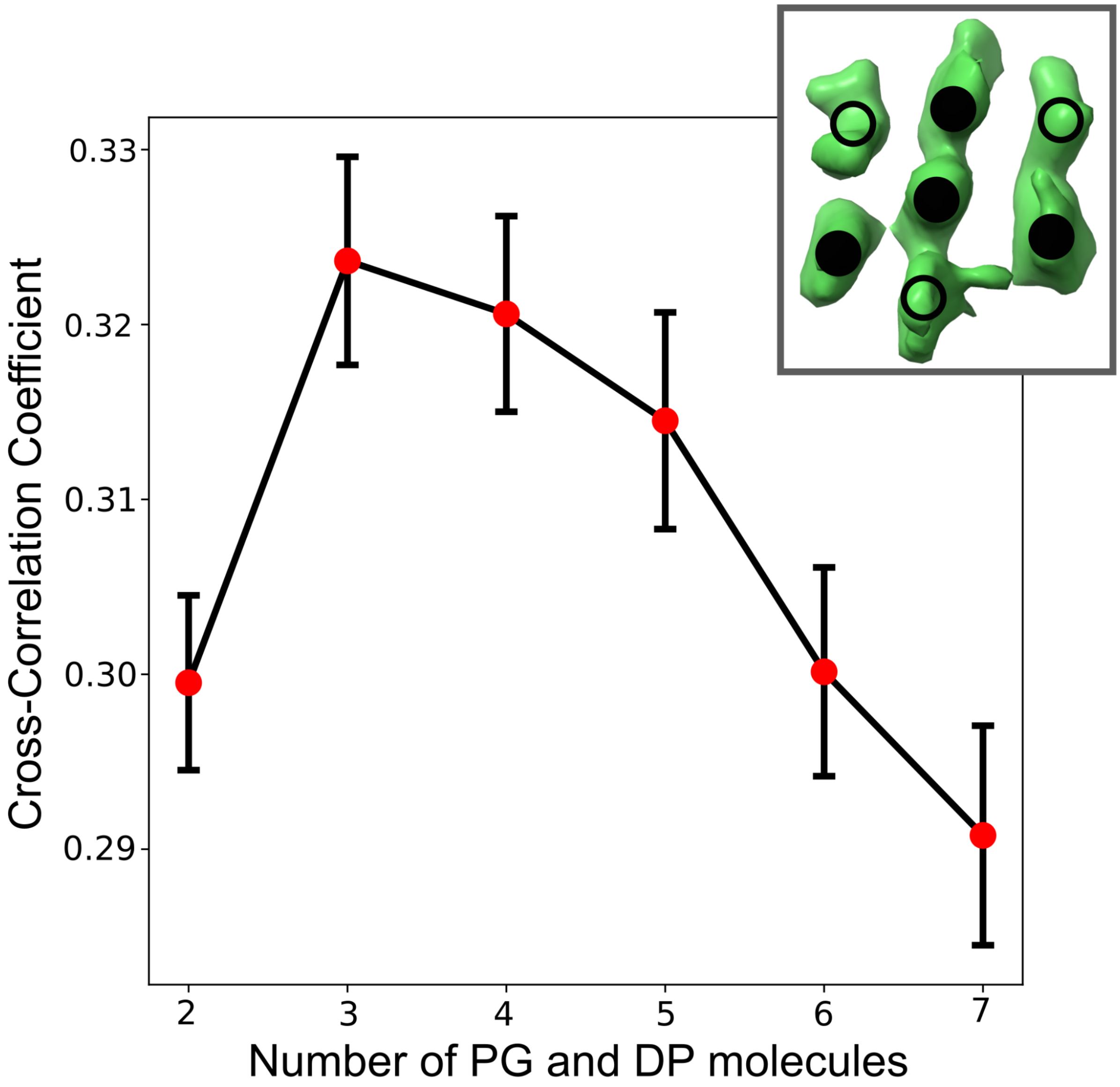


**Figure S1 Estimating the number of PG and DP copies and selecting the layout for PKP copies** The graph shows the results for independent stoichiometry runs (see Methods, Stage 2 and Supplementary Section 1.1) with the number of PG and DP molecules ranging from 2 to 7. The boxplot marks the mean (red dot) and the standard deviation (black error bars) of the cross-correlation coefficient (Bonomi et al., 2019) between the top 10% best-scoring models (based on their cross-correlation coefficient) and the cryo-tomogram in each of the runs. **(Inset)** The seven densities in the PKP layer of the tomogram (top view) of which four were full-length PKPs in our model (filled circles) and three were fixed, non-interacting PKP structured regions (empty circles).

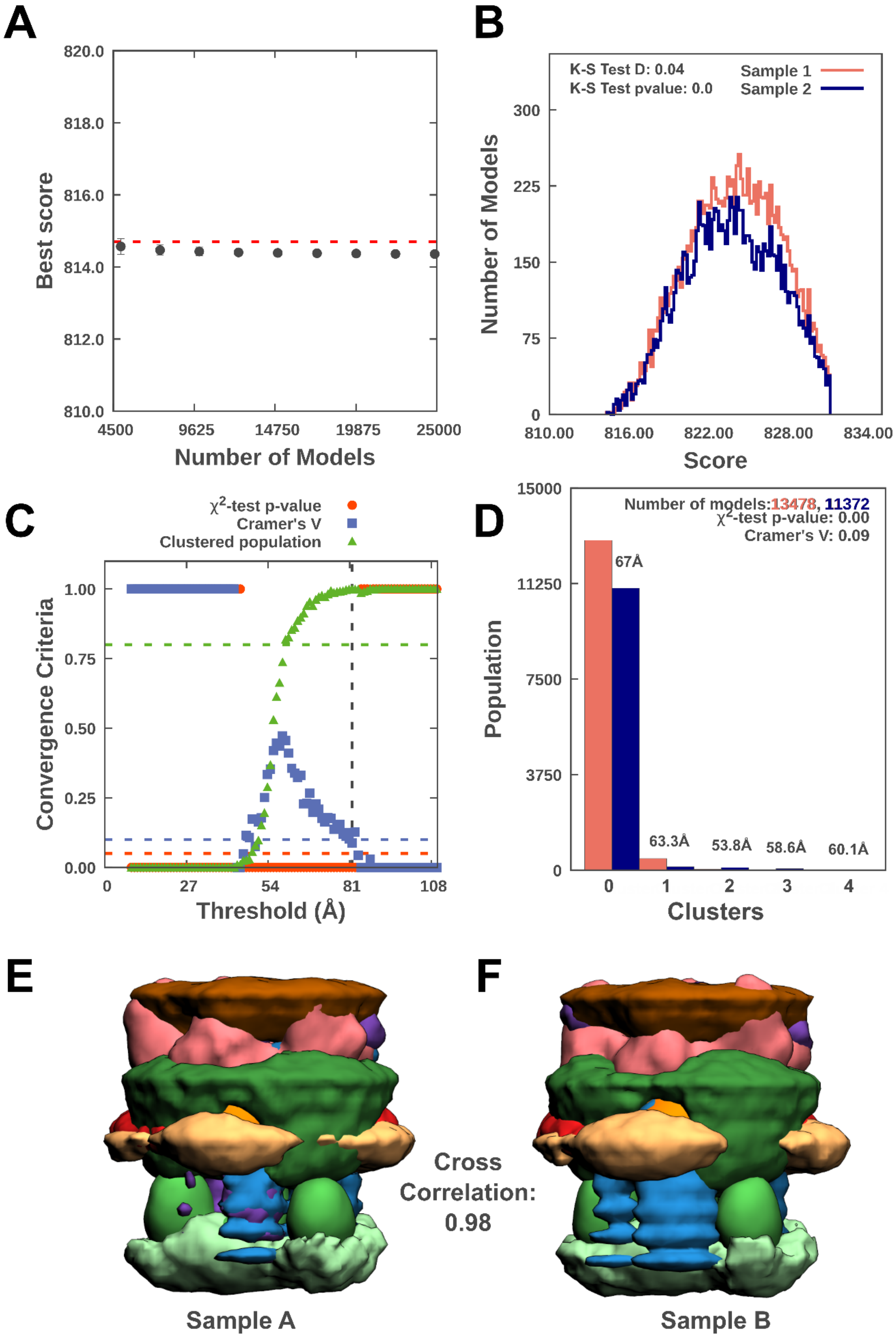

A
Best score
820.0
818.0
816.0
814.0
812.0
810.0
4500
9625
14750
19875
25000
Number of Models
B
K-S Test D: 0.04
K-S Test pvalue: 0.0
Sample 1
Sample 2
Number of Models
300
225
150
75
0
810.00
816.00
822.00
828.00
834.00
Score
C
χ2-test p-value
Cramer's V
Clustered population
Convergence Criteria
1.00
0.75
0.50
0.25
0.00
0
27
54
81
108
Threshold (Å)
D
Number of models:13478, 11372
χ2-test p-value: 0.00
Cramer's V: 0.09
Population
15000
11250
7500
3750
0
67Å
63.3Å
53.8Å
58.6Å
60.1Å
0
1
2
3
4
Clusters
E
F
Cross Correlation: 0.98
Sample A
Sample B

**Figure S2 Sampling Exhaustiveness protocol for Desmosomal ODP** (Viswanath et al., 2017)) **A)** Test for the convergence of the model score for the 24016 good-scoring models. The scores do not continue to improve as more models are added independently. The error bar represents the standard deviations of the best scores, estimated by repeating sampling of models 10 times. The red dotted line indicates a lower bound reference on the total score. **B)** Testing the similarity of model score distributions between samples 1 (red) and 2 (blue). The difference in the distribution of scores is significant (Kolmogorov-Smirnov two-sample test p-value less than 0.05) but the magnitude of the difference is small (the Kolmogorov-Smirnov two-sample test statistic D is 0.04); thus, the two score distributions are effectively equal. **C)** Three criteria for determining the sampling precision (Y-axis), evaluated as a function of the RMSD clustering threshold (X-axis). First, the p-value is computed using the $\chi^2$-test for homogeneity of proportions (red dots). Second, an effect size for the $\chi^2$-test is quantified by the Cramer's V value (blue squares). Third, the population of models in sufficiently large clusters (containing at least 10 models from each sample) is shown as green triangles. The vertical dotted grey line indicates the RMSD clustering threshold at which three conditions are satisfied (p-value > 0.05 [dotted red line], Cramer's V < 0.10 [dotted blue line], and the population of clustered models > 0.80 [dotted green line]), thus defining the sampling precision of 82 Å. **D)** Populations of sample 1 and 2 models in the clusters obtained by threshold-based clustering using the RMSD threshold of 82 Å. Cluster precision is shown for each cluster **E-F)** Comparison of localization probability densities of models from sample A and sample B for the major cluster. The cross-correlation of the density maps (see Supplementary section 2.2) of the two samples is greater than 0.98.

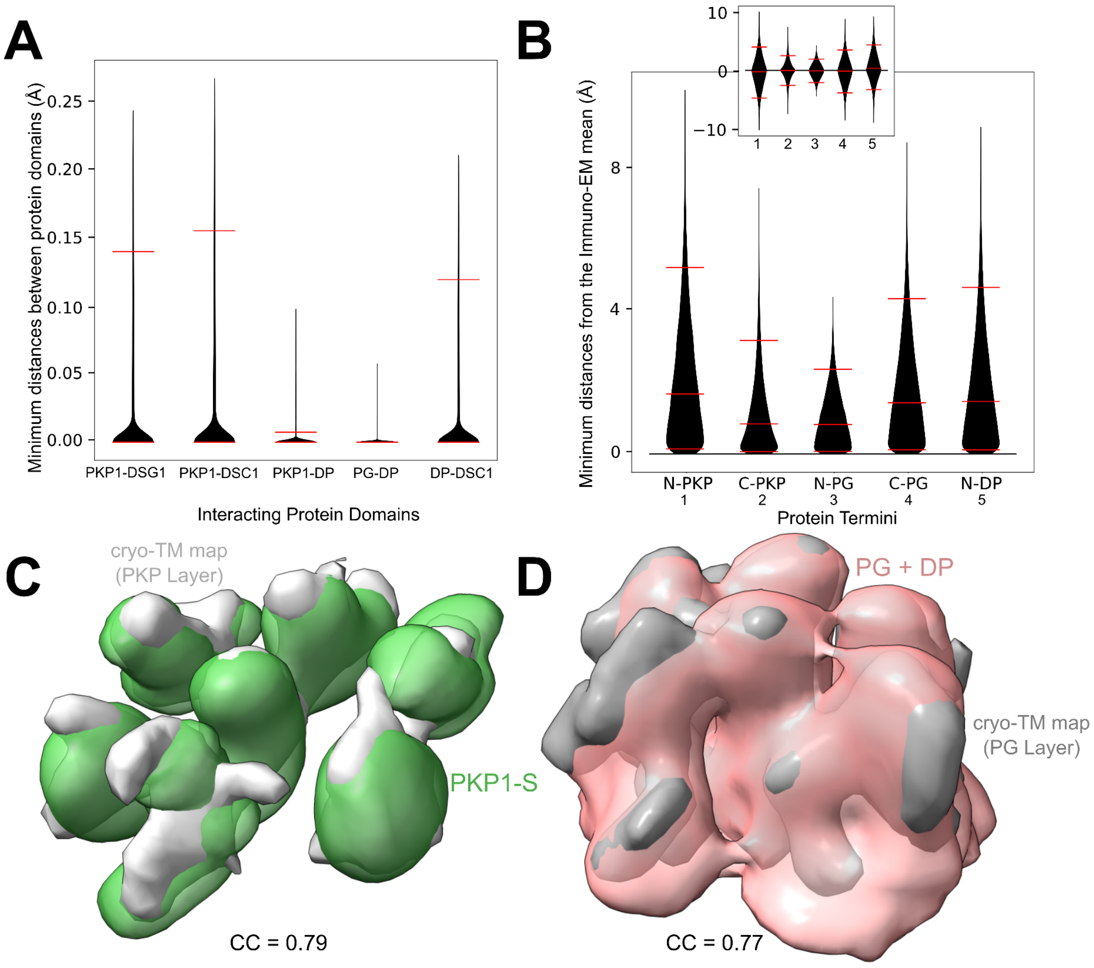


**Figure S3 Fit to data used in modeling A)** Fit to the data from biochemical experiments formulated as binding restraints (Methods). Each violin corresponds to the absolute closest distance between the two interacting domains across all protein copies for a model in the major cluster (Methods). Each distribution corresponds to a restraint in Table S2A. Red horizontal lines correspond to 5th, 50th and 95th percentile (in **A** and **B** both) after outlier removal. **B)** Fit to immuno-EM data (North et al., 1999). Each violin corresponds to the absolute difference between the experimental mean and the model-predicted distance from the membrane (Methods). The inset shows the same information without the absolute value (i.e. signed difference). **C-D)** Fit to the cryo-tomogram for the PKP Layer (**C**) and the PG Layer (**D**). Densities from the model (colored) are shown along with the segmented densities from the tomogram, EMD-1703 (Al-Amoudi et al., 2011)(Methods). The cross-correlation (CC) (Methods, Supplementary Section 2.2) is mentioned for each of the fits. PKP1-S density (including the non-interacting PKP1 molecules) and the PG + DP density are visualized at a ~10% threshold and the tomogram is visualized at the recommended threshold. See also Fig. 3, Table S2.

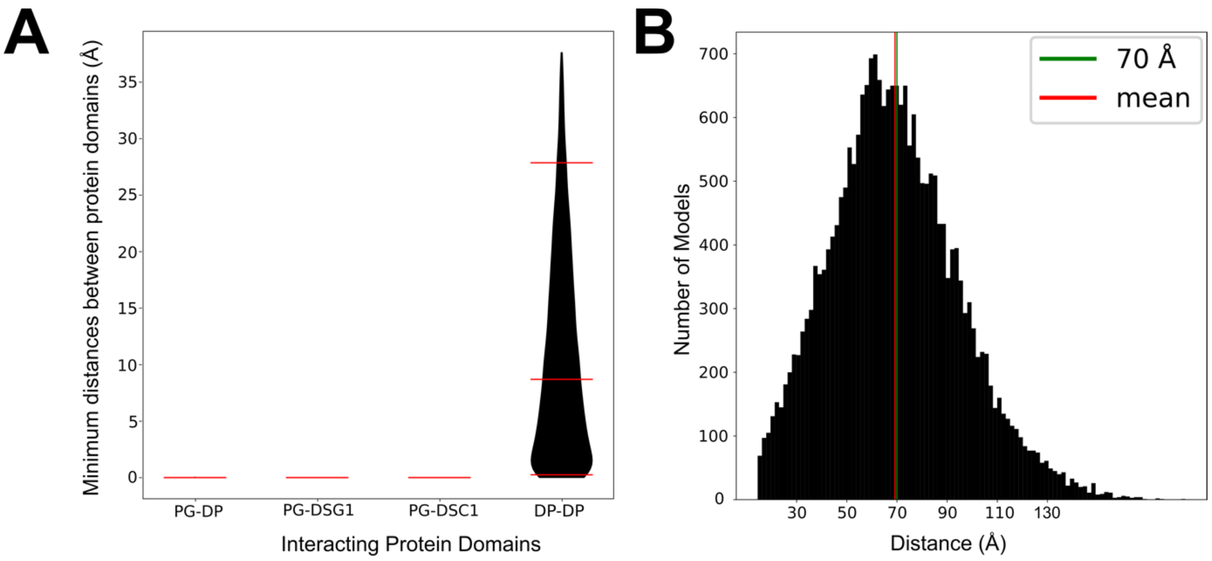


**Figure S4 Fit to data not used in modeling A)** Fit to the data from biochemical experiments not used for modeling (Methods). Each violin corresponds to the closest distance between the two interacting domains across all copies for a model in the major cluster (Methods). Each distribution corresponds to a row in Table S3. Red horizontal lines correspond to 5th, 50th and 95th percentile after outlier removal. See also Table S3. B) Fit to the cadherin spacing data from (Sikora et al., 2020). Histogram displays the minimum DSC1-DSG1 distance at the plasma membrane for each model. The data from Sikora *et*. al. is shown in green, and the mean of the distribution from integrative models is shown in red.

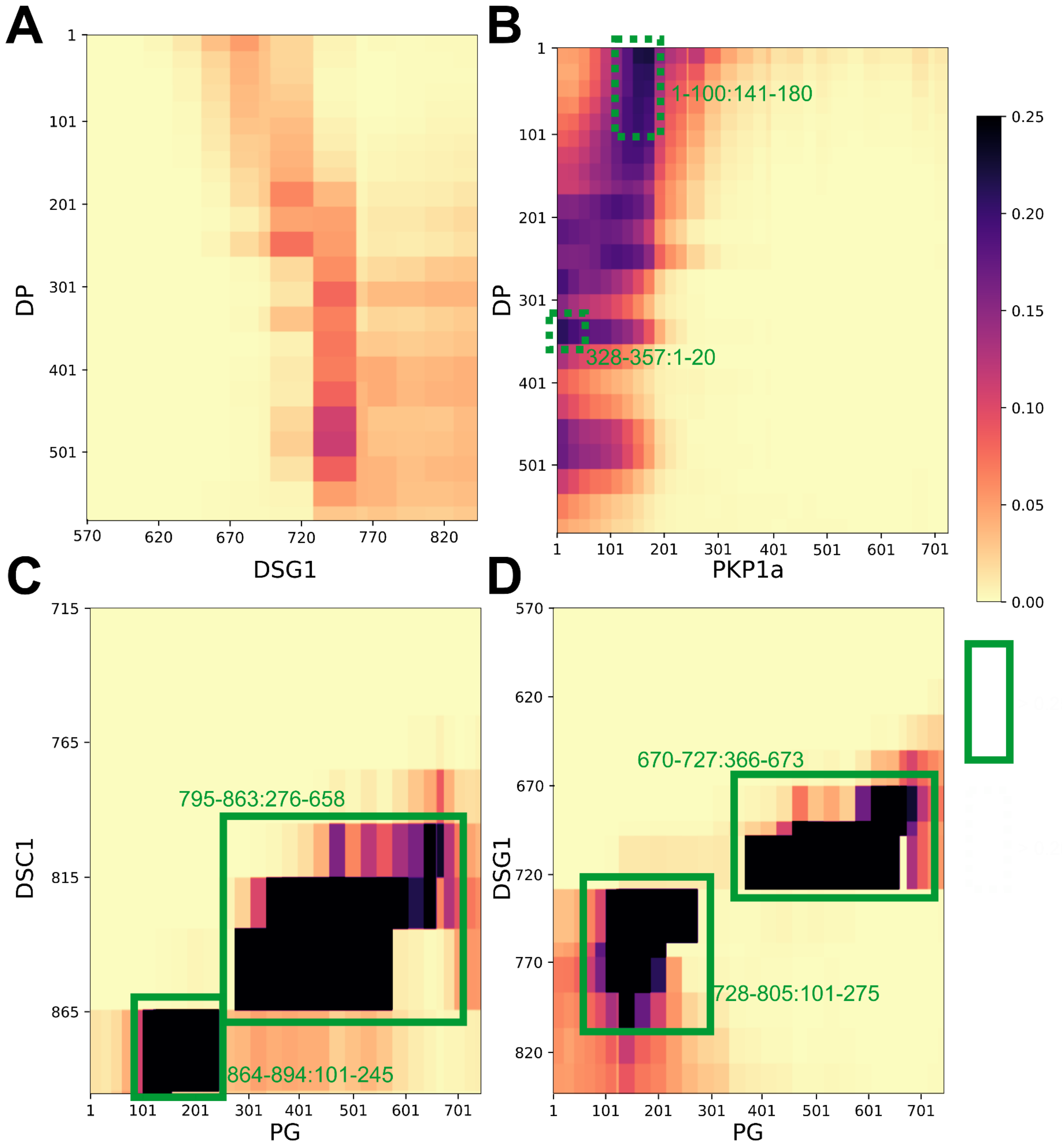


**Figure S5 Additional contact maps** Protein-protein contact maps for DP-DSG1 (**A**), DP-PKP1 (**B**), PG-DSC1 (**C**), and PG-DSG1 (**D**) pairs. Maps are colored by the proportion of the models in the major cluster where the corresponding two bead surfaces are within contact distance (10 Å). Rectangles with solid green (broken green) lines outline novel contacts present in >25% (>20%) of the models. See also Fig. 4 and Table S4.

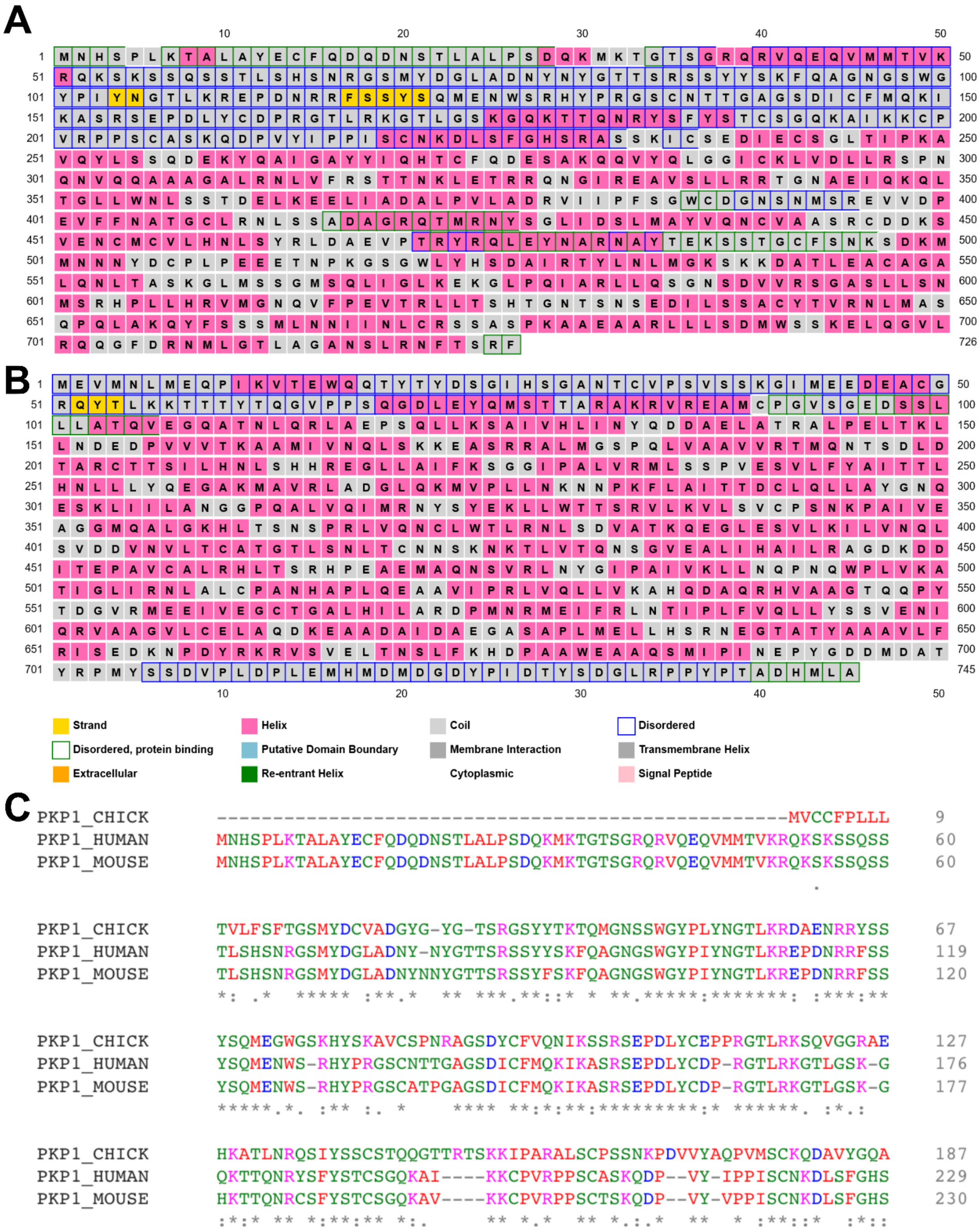


**Figure S6 Prediction of disorder in PG and PKP1 and sequence conservation for PKP1**
The PSIPRED (Buchan & Jones, 2019) output for PG (**A**) and PKP1 (**B**) are shown with the color scheme displayed below panel B. **C)** Sequence conservation output using Clustal Omega (Goujon et al., 2010; Sievers et al., 2011) for Chick, Human and Mouse PKP1$^{1\text{-}229}$. The colors represent different kinds of amino acids (acidic: blue, basic: magenta, hydrophobic/small: red,

Hydroxyl/sulfhydryl/amine/G: green). An asterisk (*) represents an exact residue match, a colon (:) represents a strongly similar match and a period (.) represents a weakly similar match.

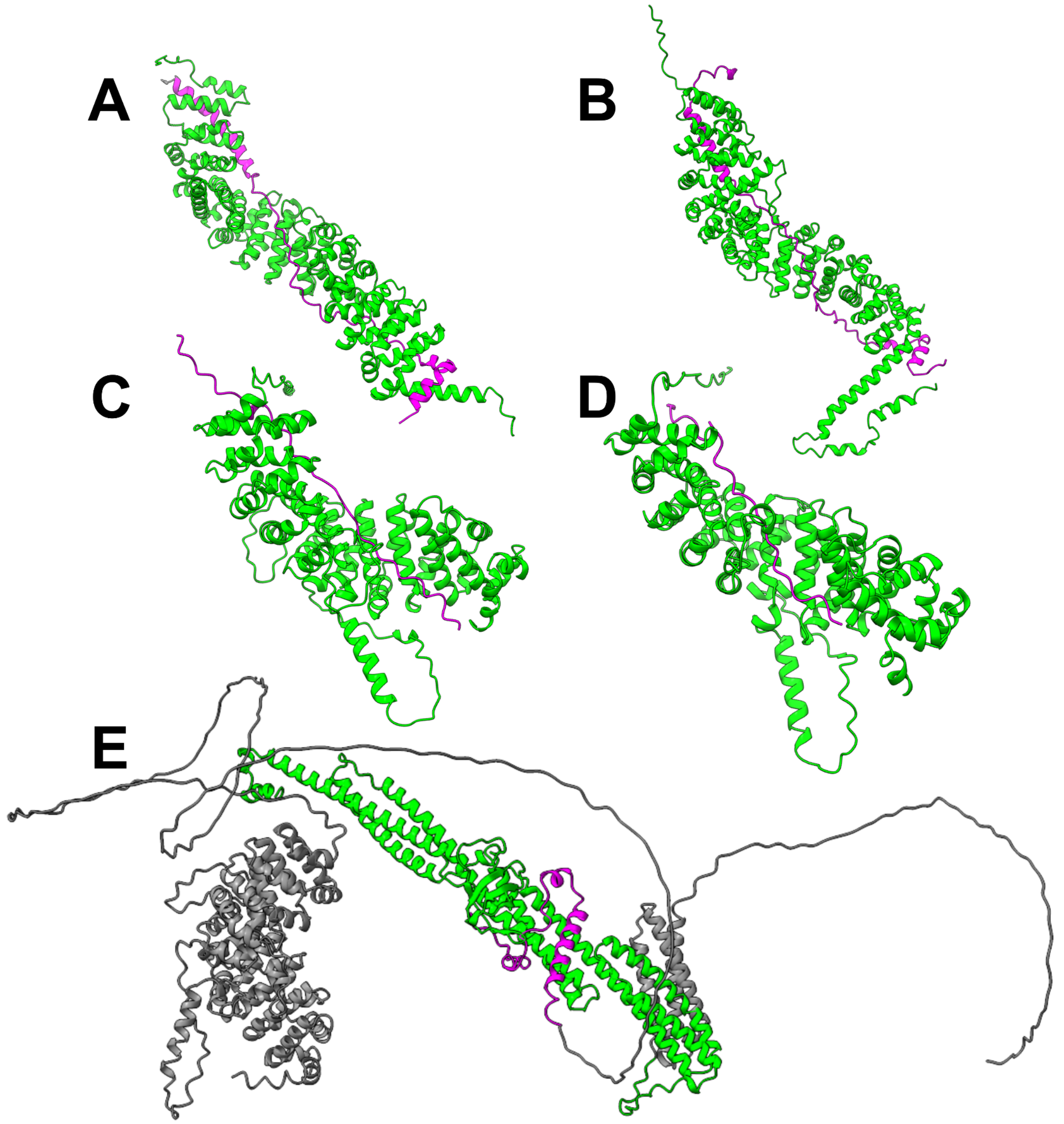


**Figure S7 Alphafold2-Multimer predictions** The best ranked model (based on PTM + IPTM score) is displayed for each protein pair. **A)** $PG^{107-697}$ (lime) and $DSC1^{812-894}$ (magenta) **B)** $PG^{75-699}$ (lime) and $DSG1^{680-768}$ (magenta) **C)** $PKP1^{229-747}$ (lime) and $DSC1^{722-759}$ (magenta) **D)** $PKP1^{230-744}$ (limer) and $DSG1^{712-737}$ (magenta) **E)** $DP^{160-584}$ (lime) and $PKP1^{1-60}$ (magenta). Rest of the residues of DP and PKP1 that are away from the interface are in gray. The PKP1 isoform used in **C, D, E** is PKP1b (in contrast to PKP1a used for integrative modeling in the paper) which has an extra 20-residue segment ($PKP1b^{412-432}$). This however does not affect any conclusions in the paper or the figure.

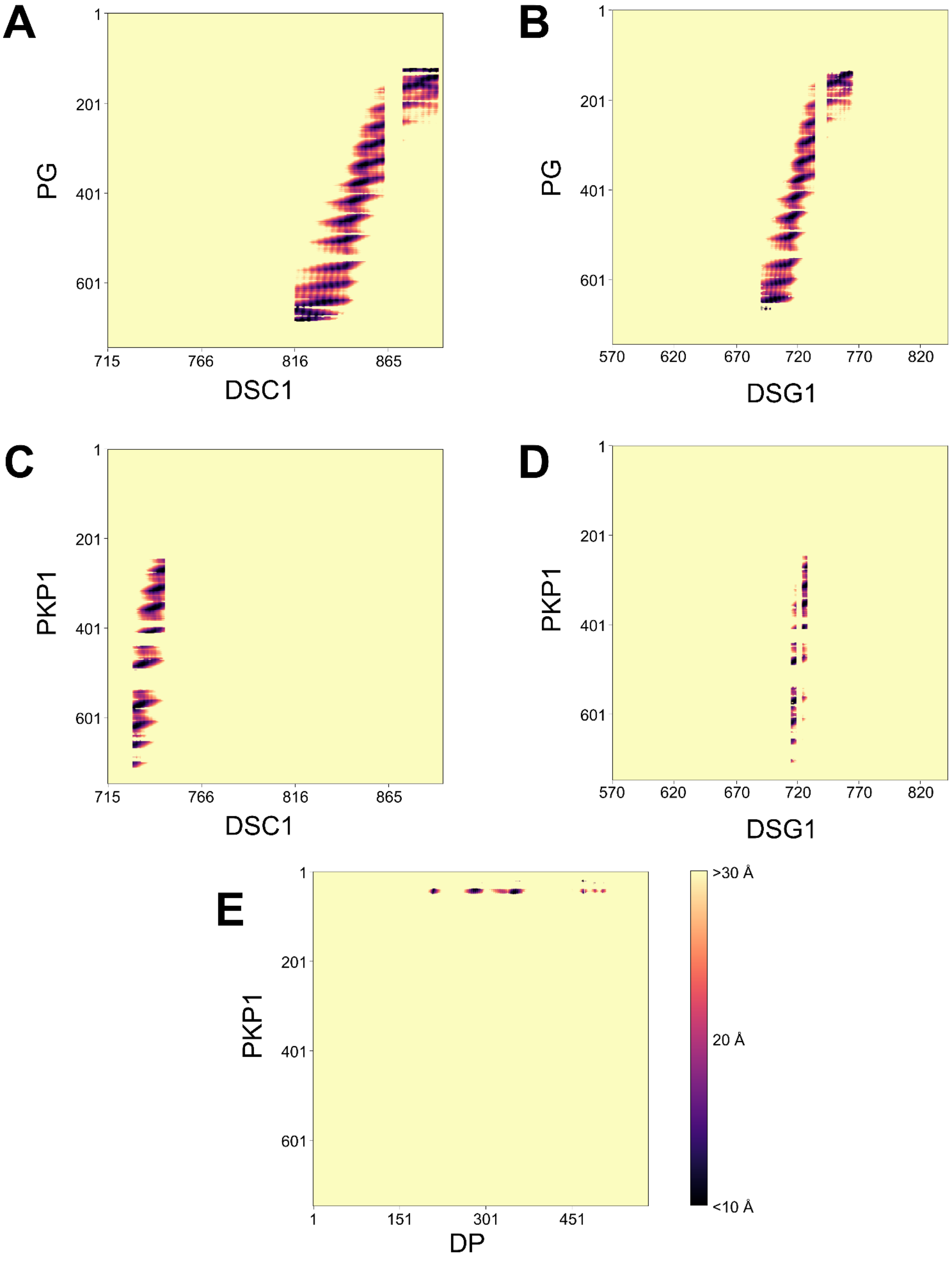


**Figure S8 Contacts predicted by Alphafold2-Multimer** Contact maps (Cα-Cα distance between residues) for PG-DSC1 (**A**), PG-DSG1 (**B**), PKP1-DSC1(**C**), PKP1-DSG1(**D**), and PKP1-DP (**E**) based on the best ranked AF2-Multimer model for each protein pair. The map is colored by the distance between Cα atoms of a residue pair. Regions in contact distance (Cα-Cα distance <10Å) are colored black. Distances are shown only for residue pairs with reliable AF2 prediction (PAE <5 and pLDDT >70 for each residue in the pair). Residue pairs without reliable AF2 prediction are colored as >30Å irrespective of their actual Cα-Cα distance. The PKP1 isoform used in **C, D, E** is PKP1b as in Fig. S8.

### Supplementary Tables

| Protein | Residue Ranges | Structure: PDB (chain) or unknown | Uniprot ID |
|---|---|---|---|
| DP1 | 1-177 | Unknown | P15924 |
| | 178-584 | 3R6N (A) | |
| PKP1a | 1-243 | Unknown | Q13835-2 |
| | 244-700 | 1XM9 (A) | |
| | 701-726 | Unknown | |
| PG | 1-125 | Unknown | P14923 |
| | 126-673 | 3IFQ (A) | |
| | 674-745 | Unknown | |
| DSG1a | 570-697 | Unknown | Q02413 |
| | 698-765 | 3IFQ (C) | |
| | 766-842 | Unknown | |
| DSC1a | 715-833 | Unknown | Q08554-1 |
| | 834-894 | 3IFQ (C) | |

**Table S1 Modeled protein domains** The different domains of the modeled proteins are shown along with their residue ranges andstructure Regions of unknown structure were represented as flexible 20-residue beads, while regions of known structure were represented as rigid bodies consisting of 30-residue beads. The colors refer to domains without a known structure (red), domains which are homology modeled on a structure template of an isoform or homolog (yellow), and domains which have the structure in the PDB (green). Only the domains in the ODP were modeled; extracellular and transmembrane domains ($DSG1^{1\text{-}569}$, $DSC1^{1\text{-}714}$) or domains outside the ODP ($DP^{585\text{-}2871}$, $DSG1^{843\text{-}1049}$) were not modeled (Al-Amoudi et al., 2011; Garrod & Chidgey, 2008; Nilles et al., 1991).

| Protein 1 | Protein 2 | Domain 1 | Domain 2 | Experiment | Reference | Data for Restraint<br>Protein 1 residues: Protein 2 residues |
|---|---|---|---|---|---|---|
| PKP1a | DSG1 | 70-213 | 570-1049 | Y2H | (Hatzfeld et al., 2000) | 70-213:570-842 |
| | | 1-286 | 568-1049 | Y2H | (Kowalczyk et al., 1999) | |
| | | 1-726 | 499-1049 | OA | (Smith & Fuchs, 1998) | |
| PKP1a | DSC1 | 1-726 | 715-894 | OA | (Smith & Fuchs, 1998) | 1-726:715-894 |
| DP | DSC1 | 1-176 | 715-894 | OA | (Smith & Fuchs, 1998) | 1-176:715-894 |
| PKP1a | DP | 1-168 | 1-584 | Y2H | (Hatzfeld et al., 2000) | 1-168:1-584 |
| | | 1-286 | 1-584 | Y2H, co-IP, Loc | (Kowalczyk et al., 1999) | |
| | | 1-726 | 1-1014 | OA | (Smith & Fuchs, 1998) | |
| | | 1-726 | 1-2871 | Loc | (Bornslaeger et al., 2001) | |
| PG | DP | 1-745 | 1-2871 | OA | (Smith & Fuchs, 1998) | 1-745:1-584 |
| **The below data are on ODP isoforms that were not included in the main model.** | | | | | | |
| PKP3a | DSC1 | 1-18 & 51-797 | 715-894 | Y2H, co-IP, Loc | (Bonné et al., 2003) | (1-18 + 51-797):715-894 |
| PKP3a | DSG1 | 1-18 & 51-293 | 519-715 Or 715-1k | Y2H, co-IP | (Bonné et al., 2003) | (1-18 + 51-293):570-842 |

| | | | | | | |
|---|---|---|---|---|---|---|
| PKP3a | DP | 19-50 | 1-2871 | Loc | (Bonné et al., 2003) | (1-18 + 51-293):1-584 & 19-50:1-63 |
| | | 1-18 & 51-293 | 1-584 | Y2H, co-IP, Loc | (Bonné et al., 2003) | |
| | | 19-50 | 1-63 | Y2H | (Bonné et al., 2003) | |
| PKP3a | DSC3 | 1-18 & 51-797 | 712-897 | Y2H, co-IP | (Bonné et al., 2003) | (1-18 + 51-797):712-897 |
| PKP3a | DSG3 | 1-18 & 51-797 | 641-999 | Y2H, co-IP | (Bonné et al., 2003) | (1-18 + 51-797):641-999 |

**Table S2A Binding Restraints** The protein-protein binding restraints are shown along with the experimental data they are based on. The restraint is formulated by including only the residues that are modeled in our ODP model. If multiple experiments provide data for a protein pair, the data from the experiment with the highest resolution is used. In Columns 3, 4 (and 5), green background (or text) represents the highest resolution information that was used to formulate the restraint and Gray represents other information that is at a lower resolution than the restraint. Experiment abbreviation are as follows: Y2H: Yeast 2 Hybrid, OA: Overlay Assay (in-vitro), Loc: Co-Localization assays, Co-IP: Co-Immunoprecipitation.

| Protein Termini | Residues | Mean dist. (Å) | SE (Å) |
|---|---|---|---|
| DP-N | 1-189 | 103 | 9.8 |
| PG-N | 1-106 | 229 | 4.5 |
| PG-C | 666-738 | 108 | 9 |
| PKP1-N | 1-285 | 158 | 11 |
| PKP1-C | 286-726 | 42 | 11 |
| **The below data are on ODP isoforms that were not included in the main model.** | | | |
| PKP3-N | 1-359 | 158 | 11 |
| PKP3-C | 360-797 | 42 | 11 |

**Table S2B Immuno-EM Restraints** The antibody binding domains for the different termini, mean distance of the termini from the plasma membrane, and the respective standard errors are shown (North et al., 1999). Domain names are in accordance with Fig. 1.

| Protein 1 | Protein 2 | Domain 1 | Domain 2 | Experiment | Reference | Data for validation<br>Protein 1 residues:<br>Protein 2 residues |
|---|---|---|---|---|---|---|
| DP | DP | 1-1014 | 1-176 | OA | (Smith & Fuchs, 1998) | 1-584:1-176 |
| PG | DP | FULL | FULL | Y2H, co-IP | (Kowalczyk et al., 1997) | 1-745:1-584 |
| PG | DSG1 | FULL | 701-768 | co-IP, Loc | (Troyanovsky, Troyanovsky, Eshkind, Krutovskikh, et al., 1994) | 123-632:701-768 |
| PG | DSG1 | 123-632 | FULL | co-IP | (Wahl et al., 1996) | |
| PG | DSG1 | FULL | 663-958 | ITC | (Choi et al., 2009) | |
| PG | DSC1 | FULL | 858-894 | co-IP, Loc | (Troyanovsky, Troyanovsky, Eshkind, Leube, et al., 1994) | 123-632: 858-894 |
| PG | DSC1 | 123-632 | FULL | co-IP | (Wahl et al., 1996) | |
| PG | DSC1 | FULL | 795-894 | ITC | (Choi et al., 2009) | |
| DP | DSC1 | FULL | 728-740 | co-IP, Loc | (Troyanovsky, Troyanovsky, Eshkind, Leube, et al., 1994) | 1-584:728-740 |
| PG | DSC1 | FULL | 728-740 | co-IP, Loc | (Troyanovsky, Troyanovsky, Eshkind, Leube, et al., 1994) | 123-632: 728-740 |

**Table S3A Validation protein-protein binding data not used in modeling** The protein-protein binding data and the corresponding references not used in modeling are shown. Data for validation was obtained using the same reasoning as in Table S2A. Experiment abbreviation are as follows: Y2H: Yeast 2 Hybrid, OA: Overlay Assay (in-vitro), Loc: Co-Localization assays, Co-IP: Co-Immunoprecipitation, ITC: Isothermal Calorimetry. Gray text represents the information that was not validated by the ensemble of models.

| Restrained protein domain (s) or residues | Data for validation | Experiment | Reference |
|---|---|---|---|
| PG-N ($PG^{1-106}$) | Distance of domain from membrane = 240 +/- 20 Å | dSTORM | (Stahley et al., 2016) |
| DP-N ($DP^{1-189}$) | Distance of domain from membrane = 410 +/60 Å | dSTORM | (Stahley et al., 2016) |
| $DSG1^{570-589}$, $DSC1^{715-734}$ | Cadherin spacing: Distance between DSG1 membrane-bound region and DSC1 membrane-bound region is 7 nm | cryoTM | (Sikora et al., 2020) |
| $DSG1^{571}$, $DSG1^{573}$ | Residues are membrane-interacting | ABE | (Roberts et al., 2016) |

**Table S3B Other validation data not used in modeling** Data from imaging and biochemistry and the corresponding references not used in modeling are shown. Experiment abbreviation are as follows: dSTORM: direct stochastic optical reconstruction microscopy, cryoTM: electron cryo-tomography, ABE: acyl biotin exchange assay. Gray text represents the information that was not validated by the ensemble of models.

| Protein 1 | Residues in protein 1 | Protein 2 | Residues in protein 2 |
|---|---|---|---|
| DP | 1-20 | DSC1 | 795-833 |
| DP | 21-60 | DSC1 | 815-833 |
| DP | 21-40 | DSC1 | 795-814 |
| DP | 61-100 | DSC1 | 815-833 |
| DP | 141-177 | DSC1 | 864-893 |
| DP | 178-237 | PG | 306-335 |
| DP | 238-267 | PG | 276-335 |
| DP | 61-80 | PG | 366-395 |
| DP | 81-100 | PG | 336-395 |
| DP | 101-140, 161-177 | PG | 306-395 |
| DP | 141-160 | PG | 306-425 |
| DP | 178-207 | PG | 246-305, 336-425 |
| DP | 208-237 | PG | 216-305, 336-395 |
| DP | 238-267 | PG | 216-275, 336-455 |
| DP | 268-297 | PG | 246-275 |
| DP | 328-357 | PG | 216-335 |
| DP | 448-507 | PG | 81-120, 126-275 |
| DP | 508-537 | PG | 61-100 |
| DP | 1-60 | PKP1 | 141-180 |
| DP | 61-100 | PKP1 | 141-160 |
| DP | 328-357 | PKP1 | 1-20 |
| DSC1 | 795-814 | PG | 636-658 |

| | | | |
|---|---|---|---|
| DSC1 | 815-833 | PG | 336-605, 636-658 |
| DSC1* | 834-863 | PG | 276-575 |
| DSC1* | 864-894 | PG | 101-245 |
| DSC1* | 894 | PG | 101-155 |
| DSC1 | 795-814 | PG | 661-673 |
| DSC1 | 815-833 | PG | 606-635 |
| DSG1 | 670-689 | PG | 606-673 |
| DSG1 | 690-697 | PG | 456-673 |
| DSG1* | 698-727 | PG | 366-658 |
| DSG1* | 728-757 | PG | 101-275 |
| DSG1* | 758-765 | PG | 101-215 |
| DSG1 | 766-785 | PG | 101-185 |
| DSG1 | 786-805 | PG | 126-155 |
| DSG1 | 670-689 | PG | 674-693 |
| DSG1* | 698-727 | PG | 659-660 |
| DSG1 | 766-785 | PG | 186-215 |
| DSC1 | 775-794 | PKP1 | 201-220 |
| DSC1 | 795-814 | PKP1 | 181-220 |
| DSC1 | 795-814 | PKP1 | 161-180 |
| DSC1 | 815-833 | PKP1 | 161-220 |
| DSG1 | 650-669 | PKP1 | 181-220 |
| DSG1 | 670-689 | PKP1 | 161-220 |
| DSG1 | 630-649 | PKP1 | 201-220 |
| DSG1 | 650-669 | PKP1 | 221-240 |
| DSG1 | 670-689 | PKP1 | 141-160 |

| | | | |
|---|---|---|---|
| DSG1 | 690-697 | PKP1 | 181-200 |

**Table S4 Protein-Protein contacts in the ODP** All the contacts (bead surface-to-surface distance of less than 10 Å) identified in at least 20% (yellow) to 25% (green) of the models in the ensemble are shown (Methods). Contacts consistent with sub-complexes of known structure are marked with an asterisk in column 1.

| Protein Domain | Residues | Mutation | Reference | Disease |
|---|---|---|---|---|
| PG-N | 19 | T → I | (Den Haan et al., 2009) | Naxos disease |
| PG-S | 265 | R → H | (Erken et al., 2011) | Naxos disease |
| PG-S | 301 | E → G | (Marino et al., 2017) | Naxos disease |
| PG-C | 680-745 | WEAAQSMIPI → GGCPEHDSHQ + Δ690-745 | (McKoy et al., 2000) | Naxos disease |
| DP-S | 287 | N → K | (Whittock et al., 2002) | Skin Fragility - Woolly Hair Syndrome |
| DP-S | 356 | T → K | (Pigors et al., 2015) | Carvajal syndrome |
| DP-S | 564 | T → I | (Boulé et al., 2012; Keller et al., 2012) | Carvajal syndrome |
| DP-S | 583 | L → P | (Keller et al., 2012) | Carvajal syndrome |
| PKP1-S | 502 | R → H | COSMIC | Cancer Mutation |
| PG-N | 4 | M → V | COSMIC | Cancer Mutation |
| DSG1 | 788 | E → K | COSMIC | Cancer Mutation |
| DSC1 | 841 | Y → F | COSMIC | Cancer Mutation |

**Table S5 Mutations** The mutations of interest in the different protein domains are shown along with the pathology associated with the mutation (Fig. 5, Results). Domain names are in accordance with Fig. 1.
